# The free amino acid-rich biostimulant, Leafamine®, promotes cell division in tomato roots and alleviates heat stress effects

**DOI:** 10.64898/2026.07.24.740532

**Authors:** Loé Malgouyre, Norbert Bollier, Pascal Martin, Marilise Nogueira, Paul D. Fraser, Nathalie Gonzalez, Emmanuelle Mounier, Michel Hernould, Frederic Delmas

## Abstract

Several tomato (*Solanum lycopersicum*) varieties are sensitive to heat stress, which compromises plant growth, development, and ultimately yield. Biostimulants represent a promising approach to improve crop performance, yet their widespread adoption is hindered by the incomplete understanding of their mechanisms of action. This study aimed to elucidate the physiological and molecular effects of a protein hydrolysate-based biostimulant (Leafamine®), in tomato, under both optimal and heat stress conditions. Leafamine® increased primary root length by 15 to 20% compared to the controls, independent of the tested conditions, through promotion of cell division and potentially expansion. Transcriptomic analyses revealed the upregulation of genes involved in cell division and expansion under both optimal and heat stress conditions and the downregulation of heat stress markers under heat stress conditions. Hormone and metabolite profiling showed elevated levels of jasmonic and salicylic acid, putrescine, citrulline, and GABA, after Leafamine® treatment, consistent with the activation of stress tolerance pathways. Leafamine® pre-treated seedlings exhibited reduced growth inhibition during heat exposure, suggesting a priming effect. These findings highlight Leafamine® as a promising biostimulant for enhancing tomato growth in the context of climate change and the potential of biostimulants generically.

## Introduction

Climate change is expected to significantly alter crop production. It is accompanied by global warming but also by exceptional climatic events triggering drought, flood, and heat stress (HS) (IPCC 2022). Current climate projections indicate a global 2°C increase in temperatures by 2041 (IPCC 2021). HS has a huge impact on the development, growth, and productivity of various crops, including tomato (Sato, Peet and Thomas, 2000; Alsamir *et al*., 2021).

HS is known to severely impair tomato (*Solanum lycopersicum*) growth and productivity by disrupting nutrient uptake, including the major nutrients such as nitrogen (N), phosphorus (P), potassium (K), iron (Fe), and boron (B) (Giri *et al*., 2017). HS also promotes the production of reactive oxygen species (ROS), such as O₂⁻ and H₂O₂, and causes oxidative damage that adversely affects cellular functions such as photosynthesis, cell division, and membrane transport (Livanos, Apostolakos and Galatis, 2012; Niu and Xiang, 2018). The heat shock factor (HSF) and heat shock protein (HSP) system plays a central role in maintaining cellular homeostasis, interacting with multiple signaling pathways involved in stress responses (Haider *et al*., 2022). Under HS, expression of most of the HSF, which is part of a large family of conserved transcription factors, is induced but at different levels and timing (Yang *et al*., 2016). HSFs are conserved transcription factors that trigger the expression of HSPs. Upon stress, HSPs act as molecular chaperones: they mitigate HS effects by preserving proteins in their functional conformations, preventing their aggregation, assisting in refolding denatured proteins, and stabilizing cellular structures (Wang *et al*., 2004; Kotak *et al*., 2007). Hormonal signaling pathways interact closely with the heat shock response pathway upon HS. For instance, in tomato plants, gibberellins (GA), salicylic acid (SA) and cytokinin (CK) mitigate oxidative damage by enhancing antioxidant enzyme activity such as superoxide dismutase (SOD), peroxidase (POD), catalase (CAT) and increasing proline accumulation, which stabilizes cellular structures (Shah Jahan *et al*., 2019; Guo *et al*., 2022; Suliman *et al*., 2024). A crosstalk between these hormones exists, allowing a multi-layer regulation of stress response and development (Verma, Ravindran and Kumar, 2016).

In tomato, it was shown that 15 days of HS, at 32°C or more, alters root biomass, root length, lateral root number, and carbon and nitrogen assimilation (Giri *et al*., 2017). By limiting nutrients and water uptake and thereby availability, root growth impairment could directly reduce fruit development and exacerbate yield losses. In Arabidopsis, this root growth reduction upon HS results from the alteration of the size and number of cells in the root apical meristem (González-García *et al*., 2023). Despite evidence of root sensitivity, the mechanisms underlying how HS affects tomato root development at the cellular and molecular level remain largely unknown, highlighting a critical gap in understanding plant adaptation to high temperatures.

Biostimulants have emerged as sustainable solution to enhance plant performance under both optimal and stressful conditions. They are described as “any substance or microorganism applied to plants that enhances nutrition efficiency, abiotic stress tolerance, and/or crop quality traits, regardless of its nutrients content” (du Jardin, 2015). Biostimulants are categorized in five main types in the literature: humic and fulvic acids, seaweed or plant extracts, amino acids (AA) and peptides, inorganic compounds including Al, Co, Na, Se, and Si, and beneficial fungi or bacteria (Yakhin *et al*., 2017). Depending on their composition, biostimulants can act either directly, by providing bioactive compounds that modulate hormonal or signaling pathways, or indirectly, by improving soil properties, nutrient availability, and rhizosphere microbial activity. Protein hydrolysate-based biostimulants (PHs) are derived from animal or plant sources and correspond to mixtures of AA, oligopeptides, or polypeptides, and the main AAs found in PHs depend on the protein source (Malécange *et al*., 2023). Several studies have reported that the application of isolated or combined AAs and PHs positively affects root growth in tomato (Colla *et al*., 2014; Ceccarelli *et al*., 2021). The promotion of root development by PHs has been associated with the modulation of nitrogen (N) assimilation in plants, by increasing the expression of amino acid transporters (*AAT1*), and repressing the expression of high-affinity nitrate transporters (*NRT2*.*1* and *NRT2.3*), which are high-affinity nitrate transporters in tomato root (Sestili *et al*., 2018). PHs have also been reported to exert hormone-like effects in tomato roots. A plant-derived PH has been described as having strong IAA-like activity, to increase the expression of genes related to GA function in plants, and to increase the enzyme activity involved in CK biosynthesis (Colla *et al*., 2014).

PHs have also been described as effective in mitigating abiotic stress in tomato roots, and their protective action involves the modulation of oxidative stress. For instance, animal-based PH enhanced tomato root development and modulated the expression of genes involved in abscisic acid (ABA) signaling, thereby mitigating the inhibitory effects of salt stress (Campobenedetto *et al*., 2021). Another animal-derived PH has been described to increase SOD, POD, and CAT activities associated with a decrease of H_2_O_2_, and O_2_^−^ under drought stress (Wang *et al*., 2022). The role of PHs in HS has been less studied. Vaseva *et al*., (2022) demonstrated that a PH could alleviate the effects of HS on maize root growth, which was associated with a decrease in *HSPs* expression. Overall, data on the impact of biostimulants on root systems under HS remain scarce, highlighting the need for further investigation in this area.

Leafamine® (LA) (BCF Life Sciences) is a PH derived from the extraction of poultry feathers and composed of 17 free L-amino acids, which make up 82% of the product, and small peptides (< 800 Da). This unusually high proportion of free amino acids distinguishes Leafamine® from many commercial PH products and may provide a rapid source of signaling molecules in addition to nutritional compounds. LA has been reported to increase leaf and root biomass, as well as protein, starch, and nitrogen contents in lettuce leaves (Malécange *et al*., 2022). Its application also elevates ABA and zeatin levels (Malécange *et al*., 2022). These results indicate a broad effect on nutritional status and hormonal homeostasis in lettuce under non-stress (NS) and drought stress (Malécange *et al*., 2022). This study aims to further investigate, using an integrative and multi-omics approach, the mechanisms underlying the effects of LA on tomato roots under NS and HS. Our findings show that LA stimulates cell division and expansion in primary roots and alleviates the stress-induced inhibition of primary root growth. Moreover, the multi-omics approach indicates that LA consistently modulates the expression of genes involved in root growth, nutrient uptake, HS response, and metabolic and hormonal pathways associated with the regulation of plant metabolic and hormonal status, thereby enhancing tolerance to HS. Moreover, the transcriptomic and metabolomic data obtained in this study suggest that LA induces a priming-like state, activating stress-response pathways even in the absence of stress.

## Material and methods

### Plant material, *in vitro* culture, and LA treatment

Tomato (*Solanum lycopersicum*) cv. Microtom were used for *in vitro* experiments. Seeds were surface sterilized in a decontamination solution containing commercial bleach (50% V/V), which represents 9.6% active chlorine) and 0.05% (V/V) Tween20® (Sigma) as surfactant. The seeds were then rinsed three times in sterile water. For each experiment, seeds were evenly sown on ¼ MS medium including vitamins (Duchefa M0222) (Murashige T & Skoog, 1962) (pH=5.8) containing 7.5 g.L⁻¹ sucrose and 4 g.L⁻¹ agar (Kalys HP696) and then, placed in a growth chamber (Hi-Point FH-740 Bionef) at 23°C/19°C day/night and a photoperiod of 16/8 h with a light intensity of 150 μmol m^-^² s^−^¹.

For the LA concentration and HS assays, 8 to 15 tomato seedlings were grown on ¼ MS medium, for 7 days before being transferred onto 24×24 cm square culture boxes (ThermoFisher Scientific 40835 245×245×25), supplemented with LA at concentrations ranging from 0.1 g.L⁻¹ to 6.4 g.L⁻¹. Glutamine (0.146 g.L⁻¹) was used as a control for free amino acid-based source of nitrogen equivalent to LA (0.2 g.L⁻¹) nitrogen content. The same environmental conditions used for germination, described above, were used for seedlings growth in NS, whereas HS consisted of 33°C during the day and 29°C at night for 6 days (Figure 1A). Tomato seedlings were grown for 7 days, and root length was recorded daily by marking root tips to monitor growth according to Figure S1A.

**Figure 1.**
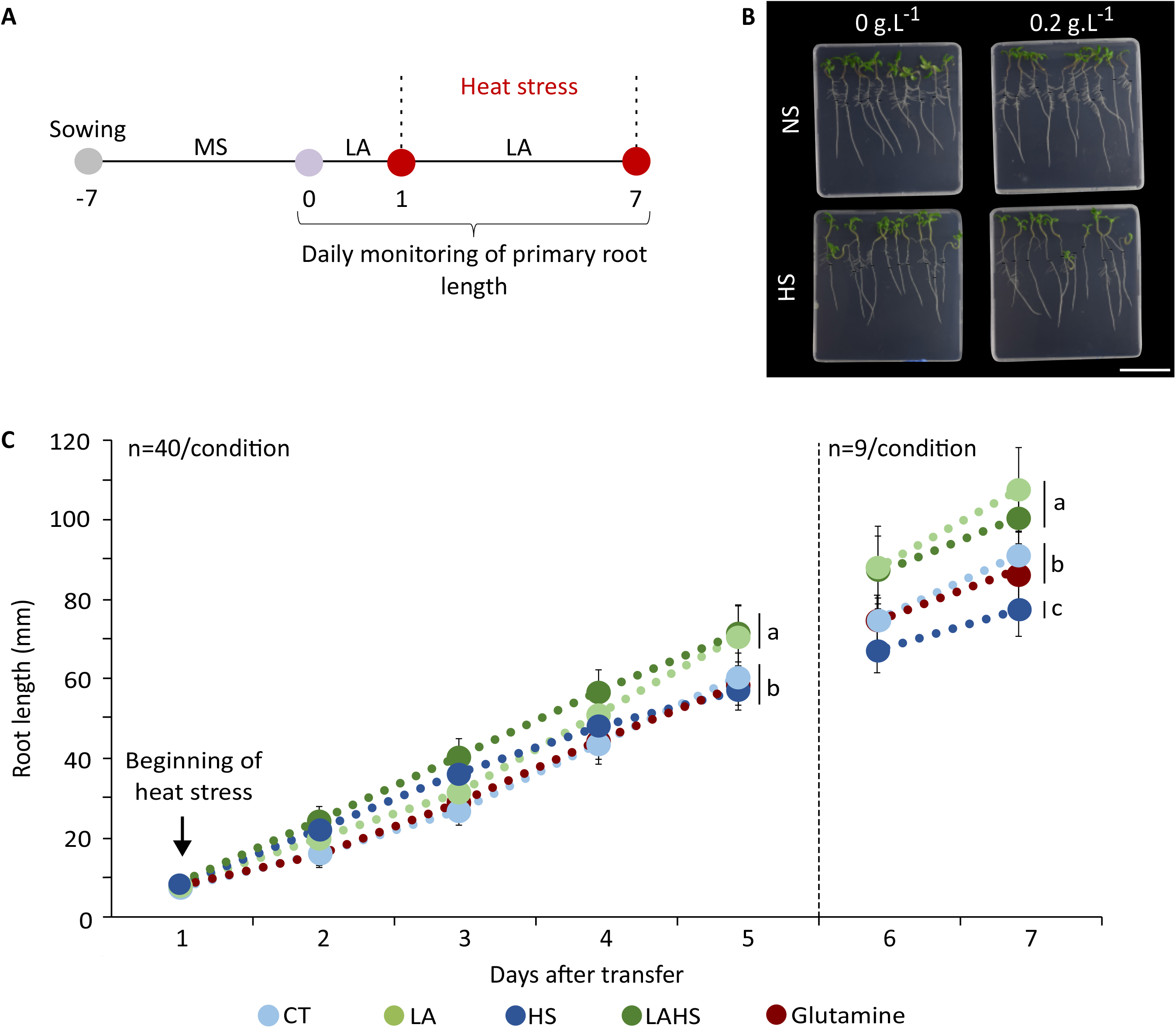
Effect of LA treatment (0.2 g.L⁻¹) on primary root growth of tomato seedlings grown under non-stress (NS) and heat stress (HS) (33 °C) conditions. (A) Experimental design of the Heat Stress (HS) assay in combination with Leafamine (LA) treatment. (B) Representative image of the seedlings grown on petri dishes under NS conditions or exposed to HS for seven days on 1/4 Murashige and Skoog (1962) media with or without LA. Scale bar = 8 cm. (C) Kinetics of Primary root growth under NS and HS conditions with or without LA treatment. Between days 1 and 5, n=40 were monitored per condition; from day 5, n=9 seedlings per condition. ANOVA test was used followed by Tukey’s multiple comparisons test. Letters indicate significant differences among treatments. Statistical comparisons (p-value) are provided in Table S2.

For the priming effect experiments, seedlings were grown on the sowing medium for 3 days before transfer into 11×11 cm square petri dishes (Greiner Bio-One 688102) with/without LA. Tomato seedlings were grown for 5 days under NS. Then, they were transferred in ¼ MS containing 7.5 g.L⁻¹ sucrose and 8 g.L⁻¹ agar medium without LA into 24×24 cm square culture boxes and grown under HS (as described above) for 7 days (Figure 7A).

LA used in these assays was dissolved in mQ water and then sterilized by autoclaving at 120°C for 20 minutes, and glutamine was filtered through a 0.22 µm PVDF filter (Millipore) and added to ¼ MS medium.

### Morphometric phenotyping

For root measurement, the primary root length was monitored daily. Each experiment was conducted at least three times, using 5 plates with 8 seedlings per plate up to day 5 and 9 seedlings from 2 plates on day 6-7 for root tracking. The fresh mass of the aerial parts was measured in groups of 12 to 22 seedlings, 5 days after transfer from the sowing medium to the treatment conditions. A total of 10 groups of seedlings under NS and 6 groups of seedlings under HS were used for the aerial fresh mass measurements. Dry mass was measured after placing the samples in a 60°C drying oven for 2 days.

### Confocal microscopy analyses of root tips

Tomato plants (*Solanum lycopersicum*) cv. Microtom *35S::NLS-GUS-GFP* lines were used for this experiment (Fernandez *et al*., 2009). Root tips, 2-cm long, were sampled 5 days after their transfer onto 24×24 cm square culture boxes with/without LA medium. Twelve root tips (3 technical repetitions) per condition were analyzed. Roots were fixed with 3.7% (V/V) formaldehyde (Formaldehyde 36% 20909.290 VWR Chemicals), vacuum-infiltrated twice for 15 minutes each and rinsed with 1X PBS (8 g.L⁻¹ NaCl, 0.2 g.L⁻¹ KCl, 1.44 g.L⁻¹ Na2HPO4, 0.24 g.L⁻¹ K2HPO4). Samples were then transferred into ClearSee solution (Kurihara *et al*., 2015) (25% (w/v) urea, 10% (w/v) Xylitol, and 15% (w/v) sodium deoxycholate) for three weeks. Then, the root tips were stained with Calcofluor white M2R (F3543, Sigma-Aldrich) at 100 µg.mL⁻¹ for 24 hours, mounted between the slide and coverslip in ClearSee and luted with nail polish. Imaging was performed using a Zeiss LSM 880 confocal microscope with a Zeiss C PL APO x63 oil-immersion objective (numerical aperture 1.4). For calcofluor imaging, excitation was performed at 405 nm and fluorescence emission collected between 405 and 438nm and for GFP, excitation was performed at 400nm and emission collected between 490 and 525 nm. The division zone was defined as the distance between the Quiescent Center (QC) and the root zone where the epidermal cell length double. For the measurement of the elongation zone corresponding to the zone comprised between the division zone and the differentiation zone where the root hairs develop, pictures of root tips were taken with a Zeiss Axioimager M2 microscope equipped with a Neofluar 10x dry objective (numerical aperture 0.3) and a Cool Snap HQ2 CCD Camera, and the distance from the root tip to the first root hair was measured using Fiji. The length of the elongation zone was then calculated by subtracting the length of the division zone from this measurement.

### RNA extraction

Roots and aerial parts were sampled 1, 3, 5, and 7 days after their transfer onto 24×24 cm square culture boxes on medium with/without LA. For each condition, three samples, corresponding to two-cm long root tips, from 3 experimental repeats were collected and immediately flash-frozen in liquid nitrogen. For total RNA extraction, 1 mL of TRIzol™ Reagent (Thermofisher Scientific) was added to the samples and then thawed at room temperature for a few seconds before grinding the samples with glass beads on an MP Fast-prep-24 (45 sec, three times, power 6). Then, 200µl of chloroform was added, samples were manually shaken for 15s, and incubated at room temperature for a few minutes. After centrifugation at 4°C for 30 min, the supernatant was collected and RNAs were purified using the RNeasy Kit (Qiagen) according to the manufacturer’s instructions RNase-free DNase (Qiagen) treatment was performed on each sample.

### RNA-seq analysis

Total RNAs were extracted as described for RT-qPCR. Libraries preparation and 2×150bp paired end sequencing by DNA Nanoball sequencing (DNBseq) was performed at the Beijing Genome Institute (BGI) on a G400 sequencer. After quality control with FastQC, reads were mapped to the SL4.0 tomato reference genome (solgenomics.net) using STAR/2.7.9a (Dobin *et al*., 2013). Pairs with unmapped reads or a mapping quality lower than 10 were filtered out. For the assignment of reads to gene models, we used featureCounts (v2.0.1) using Solanum lycopersicum ITAG4.1 annotation. The global sample topology was evaluated by principal component analysis with the FactoMineR (v2.10) R/Bioconductor package (Lê, Josse and Husson, 2008) based on the 500 most variable genes. First, we identified genes with a significant interaction between LA and HS using the Likelihood Ratio Test (BH-adjusted p-value < 0.05, Table S3) in DESeq2 (v1.42.1) R/Bioconductor package. Among these, we identified genes with similar expression patterns by clustering them by hierarchical clustering after rlog normalization using 1-Pearson correlation coefficient as distance and Ward.D2 as agglomeration method. To focus on specific comparisons, we then applied a model with a single group factor, calculated statistics for the contrasts LA vs CT (Table S7) and LAHS vs CTHS (Table S9) and selected genes with a BH-adjusted p-value < 0,05 and an absolute log2 fold change > 0.5. Enrichment of Gene Ontology Biological Process annotations were performed with the clusterProfiler (v4.10.1) R/Bioconductor package (Wu *et al*., 2021) and Plaza 5.0 annotations (https://doi.org/10.1093/nar/gkab1024).

### Extraction and analysis of metabolites and phytohormones

Roots of 6 biological replicates per condition, representing 40 seedlings per sample, were sampled 5 days after their transfer on medium with/without LA. Samples were lyophilized before being ground with metal beads. Three extractions were performed on each biological replicate (10 mg of freeze dried powder). Extraction of a quality control (mix of powder from all samples) and a blank extraction (no sample powder) were also performed with each extraction batch.

Extraction and analysis of metabolites were performed as described in (Nogueira *et al*., 2024). Respectively, deuterated succinic acid and deuterated myristic acid were used as internal standard in the separated polar and non-polar phases.

Twenty mg of freeze-dried powder were used for the phytohormones analysis. Extraction and analysis were performed exactly as described in (Nogueira *et al*., 2024).

### Statistical analysis

Statistical analyses of data other than RNA-seq were conducted using GraphPad Prism 9.0.0 for Windows. Statistical details are available in the figure legends. One-way ANOVA followed by Tukey’s multiple comparison test or Kruskal-Wallis test followed by Dunn’s test were applied depending on the normality of the data analyzed with the Shapiro-Wilks test.

## Results

### LA stimulates tomato primary root development under standard and high ambient temperatures

We evaluated the potential dose-effect relationship of LA on tomato primary root growth by monitoring root length daily after the transfer of seven-day-old seedlings onto media supplemented with LA at concentrations ranging from 0.1 to 6.2 g.L⁻¹ (Figure S1). While the concentration of 0.2 g.L⁻¹ significantly promoted root growth, concentrations of 0.1 g.L⁻¹, 0.4 g.L⁻¹, and 0.8 g.L⁻¹ did not induce significant changes. Higher concentrations, over 0.8 g.L⁻¹, resulted in a reduction of root growth (Figure S1, Table S1). The optimal LA concentration of 0.2 g.L⁻¹ was thus retained for subsequent experiments.

To evaluate LA effects on root thermotolerance, seedlings were treated or not with 0.2 g.L⁻¹ LA and exposed to 25°C (NS) or 33°C (HS) for seven days (Figure 1A). At day 7, under NS, LA-treated seedlings (LA) showed a 18% significant increase in root length compared to untreated and unstressed control seedlings (CT). HS alone significantly reduced root length by 15% after five days, whereas LA-treated seedlings under heat stress (LAHS) developed roots 29% significantly longer than untreated stressed seedlings (HS) and thus with similar length as CT seedlings at day 7 (Figure 1B, C). These results demonstrate that LA promotes root growth under NS and mitigates HS-induced growth inhibition. However, LA had no significant effect on shoot fresh nor dry biomass under neither NS nor HS conditions (Figure S2).

LA contains AA and could potentially influence root growth through nitrogen supply rather than through a signaling effect. To distinguish potential signaling effects of LA from any potential effect of a nitrogen supply by the AA mix, seedlings were grown on medium supplemented with glutamine at a nitrogen concentration equivalent to 0.2 g.L⁻¹ LA. Glutamine treatment did not induce significant changes of the mean root length compared to CT, suggesting that the LA growth-promoting effect is due to its specific composition rather than its nitrogen content alone triggering a signaling effect (Figure 1C, Table S2).

Together, these findings indicate that LA promotes root growth under NS and mitigates the inhibitory effects of HS on root development without altering shoot biomass under *in vitro* culture conditions.

### LA affects cell number in the primary root meristem

To investigate the cellular effects of LA on the root, LA-treated root tips organization were compared to CT root tips under NS or HS conditions (Figure 2A). Root tips were sampled five days after transfer and stained with calcofluor to visualize cell walls, allowing precise delineation of the main developmental zones: the columella, the division zone (root meristem), and the elongation zone.

**Figure 2.**
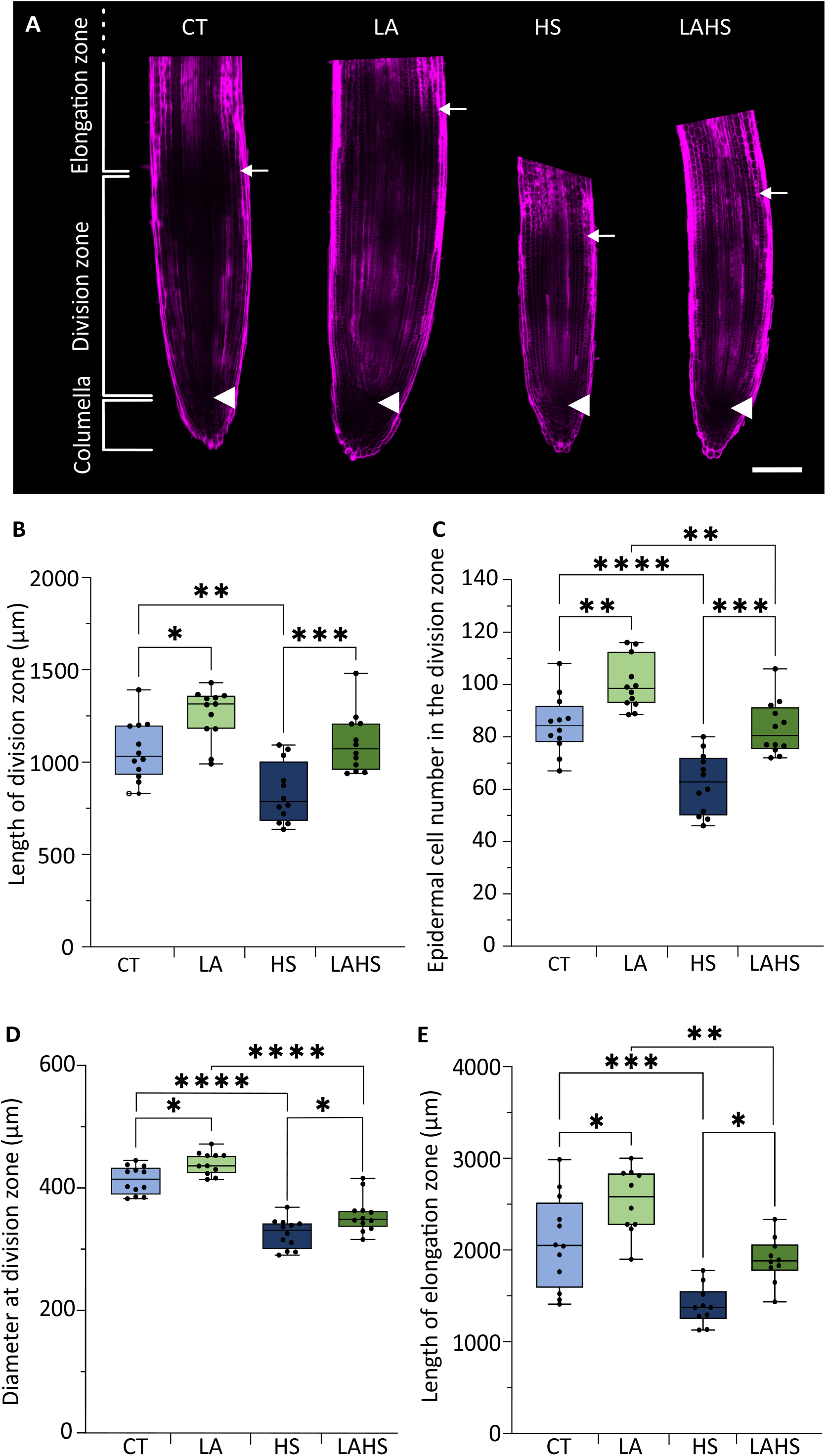
LA treatment stimulates cell divisions and cell expansion in root apical tip under NS and HS conditions. (A) Confocal images of primary root apical meristems stained with calcofluor. Arrows delimit the upper boundaries of the division zone, and arrowheads indicate the lower boundaries at the Quiescent Center (QC). Scale bar = 200µm. (B) Length of the division zone. (C) Epidermal cell number within the division zone. (D) Root diameter at the division zone. (E) Length of the elongation zone. Boxplot: whiskers extend from minimum to maximum, box extends 25th to 75th percentiles, the line in the middle is the median. Asterisks indicate significant differences (ANOVA followed by Tukey’s multiple comparisons test); *p < 0.05, **p < 0.005, ***p < 0.0005, ****p < 0.0001.

Under NS, LA significantly increased the length of the division zone by 18% compared to CT, while HS alone significantly reduced it by 21% compared to CT. Under HS, LA treatment (LAHS) resulted in the maintenance of a division zone length comparable to CT, with no significant difference between them, while being 32% significantly longer than HS seedlings (Figure 2B).

To determine whether the increased length of the division zone resulted from enhanced cell proliferation, the number of epidermal cells along the meristem longitudinal axis was measured. LA significantly increased epidermal cell number by 18% under NS, while HS significantly reduced it by 26%. Notably, root tips of LAHS seedlings recovered this deficit, showing 33% more cells than HS seedlings (Figure 2C). LA also resulted in a significant increase in root diameter by 7% compared with CT seedlings, whereas HS significantly reduced root diameter by 20% in CT, but LAHS seedlings were significantly 9% wider than HS seedlings (Figure 2D).

To further characterize the cellular effect of LA, the length of the elongation zone in the primary root was quantified. Under NS, LA significantly increased the length of this zone by 33% compared to CT. HS significantly reduced the elongation zone length by 32% relative to CT; however, in LAHS, the elongation zone was 35% significantly longer than in HS (Figure 2E; Figure S3).

In conclusion, LA restored division zone cell number and elongation zone length in HS seedlings to levels comparable to CT in NS, suggesting that LA promotes cell division under NS while preserving meristem activity and potentially sustaining cell expansion under HS.

### LA and HS modulate the expression of genes related to protein homeostasis

To investigate the molecular effects of LA on roots under NS and HS, genome-wide transcriptomic analysis was performed. Root tip samples were collected five days after transfer to LA-supplemented media, corresponding to four days after exposure to HS for the stressed plants, as illustrated in Figure 1C. Total RNA was extracted and subjected to RNA sequencing.

Across the entire dataset, 26,627 expressed genes were detected with at least one count for one sample. Principal component analysis (PCA) of the 500 most variable genes showed that the expression profile of CT, HS, LA and LAHS samples were clearly distinct as they all clustered independently (Figure 3A). The HS caused the greatest variation (Dimension 1: 54.26% of the total variance) followed by the LA (Dimension 2: 23.26% of the total variance). This indicates that both treatments (LA and HS) significantly influenced gene expression.

**Figure 3.**
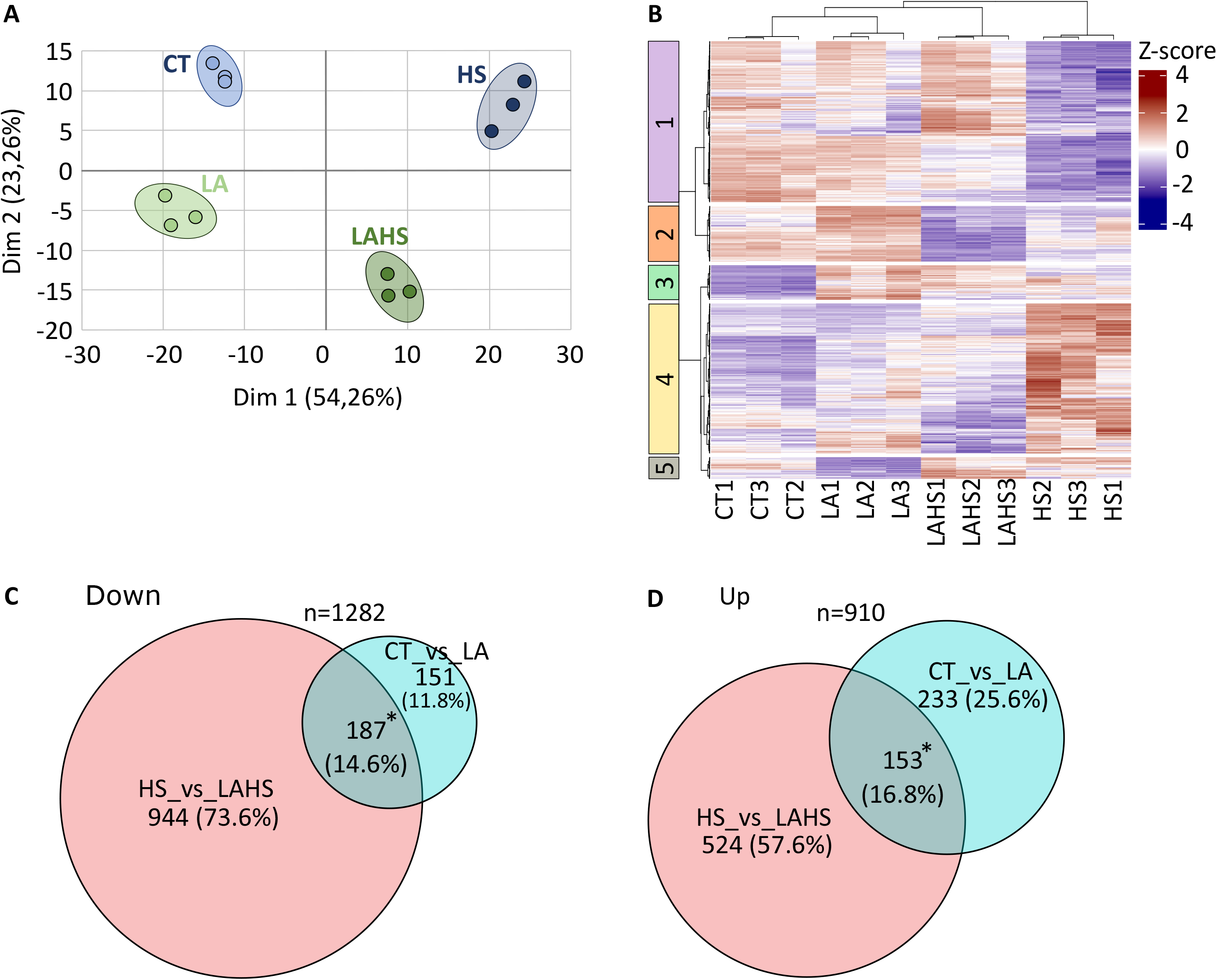
LA treatment modifies the transcriptomic responses of tomato roots under NS and HS conditions. (A) Principal component analysis (PCA) using the 500 most variable genes identified. (B) Expression pattern of DEGs clustered according to the interaction between LA treatment and HS. Normalized gene expression values (z-scores) are represented using the indicated colour scale. (Log2FC > 0.5). (C) Venn diagram showing the DEGs upregulated by LA treatment (Log2FC > 0.5) identified under NS conditions, HS, or both. (D) Venn diagram showing DEGs downregulated by LA treatment (Log2FC < 0.5) identified under NS conditions, HS conditions, or both. Asterisks indicate significant differences (Hypergeometric test); *p < 0.05.

To evaluate the combined effects of both LA and HS on gene expression, we searched for genes with a significant treatment:stress interaction (DESeq2, LRT, BH-adjusted p-value < 0.05). This approach identified 761 differentially expressed genes (DEG) whose expression response to stress was dependent on LA treatment (Table S3). These DEGs were grouped into five clusters by hierarchical clustering (Figure 3B, Table S3).

Gene Ontology (GO) enrichment analysis was performed for each cluster and revealed significant terms (*p* < 0.05) only in clusters 2 and 4 (Table S4). All GO terms were manually grouped to remove redundancy (Figure S4A). Genes in cluster 2 (100 genes) showed higher expression in LA-treated roots under NS, but lower in LAHS roots. This cluster was enriched for 57 GO terms, which were related to stress response (including HS), and to protein metabolism (proteolysis, protein folding, proteasomal protein catabolic process, ubiquitin-dependent protein catabolic process) (Figure S4A). In cluster 4 (270 genes), transcripts were highly expressed in HS samples compared to other conditions, with 32 GO terms significantly enriched (Figure S4A). Many terms were related to protein metabolism and primary metabolism (e.g., Cellular metabolic process, primary metabolic process, macromolecule metabolic process, regulation of macromolecule biosynthetic process) (Figure S4A). Although no GO terms reached statistical significance in cluster 1, three terms related to root development: trichoblast differentiation, root epidermal and cell differentiation, were found associated with this cluster (Figure S4A).

In summary, this analysis indicated that genes associated with protein folding and HS response are downregulated by LA under HS, while being highly expressed after LA treatment under NS, suggesting that LA may prime roots for improved stress resilience. Priming is defined as a “mechanism which leads to a physiological state that enables seedlings to respond more rapidly and/or more robustly after exposure to biotic or abiotic stress” (Aranega-Bou *et al*., 2014).

### LA stimulates the expression of cell division markers and modulates the expression of HS markers

To better understand the mode of action of LA, the pathways modulated by LA under both NS and HS conditions were analyzed. DEGs (adjusted *p* < 0.05 and Log2FC > 0.5) were identified by conducting two pairwise comparisons under NS (LA versus CT) and HS (LAHS versus HS), yielding 723 DEGs and 1808 DEGs, respectively. Under NS, LA triggered the downregulation of 338 genes, whereas under HS, 1,131 genes were downregulated, with 187 genes common to both conditions (Figure 3C, Table S5). Similarly, LA induced the upregulation of 386 genes under NS and 677 under HS, with 153 genes commonly upregulated (Figure 3D, Table S5).

GO BP enrichment analysis of genes commonly regulated by LA revealed 28 enriched terms in the list of downregulated (187 genes) genes and 75 enriched terms in the list of upregulated genes (153 genes) (Table S6). All GO terms were manually grouped to remove redundancy, as for Figure S4B. GO terms among the commonly downregulated genes were associated with stress response, metabolism, transport, and the cell wall process, while GO terms of the upregulated genes were linked to biotic stress, SA metabolic process, specialized biosynthetic process, different types of transports, and cellular response to starvation, suggesting that LA may influence nutrient status in roots (Figure S4B, Table S6).

Next, we focused on genes specifically regulated by LA treatment under NS (Table S7). A GO enrichment analysis of the 338 downregulated genes identified 27 significantly enriched GO terms associated with cell wall organization, response to nitrogen compounds, antioxidant activity, and detoxification (Figure 4A, Table S8).

**Figure 4.**
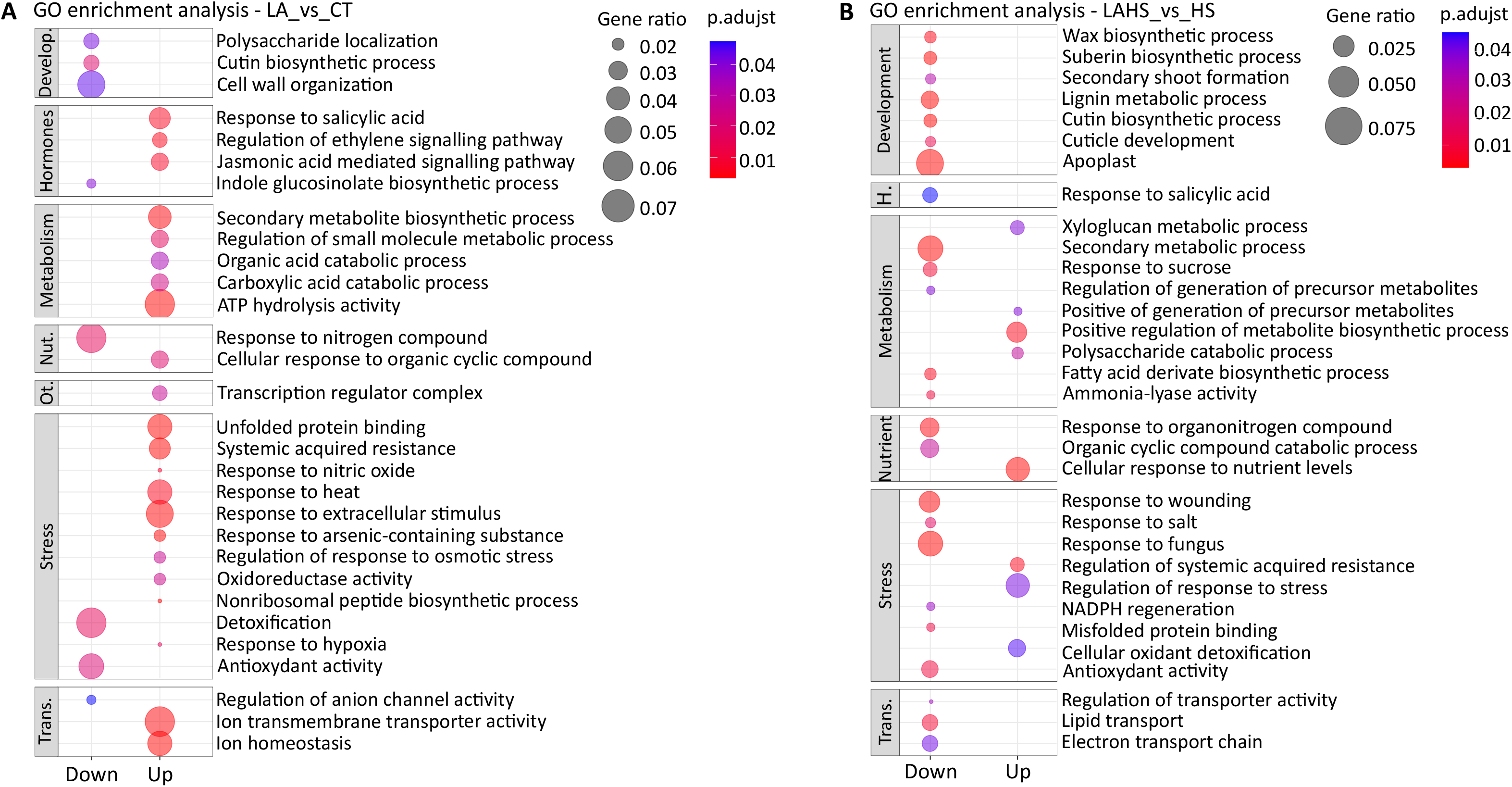
LA treatment depending on environmental conditions specifically impacts cellular and molecular processes associated to development, metabolism and stress response (A) GO enrichment analysis of DEGs (Log2FC > 0.5, Log2FC < -0.5), both up-and downregulated by LA treatment under NS conditions. (B) GO enrichment analysis of DEGs (Log2FC > 0.5, Log2FC < -0.5), both up-and downregulated by LA treatment under HS conditions. Develop = Development. H = Hormones. Nu = Nutrition. Ot = Others. Trans = Transcription. In GO analyses, terms are grouped by functional categories. Each dot represents a GO term, coloured according to the adjusted p-value and sized according to the gene ratio.

Among the 386 upregulated genes, 150 GO terms were significantly enriched (Figure 4A, Table S8). Enriched terms included hormone signaling pathways (response to SA, regulation of ethylene-activated signaling pathway, and jasmonic acid (JA) mediated signaling pathway), specialized metabolite biosynthetic process, response to abiotic stress (response to heat, response to extracellular stimulus, regulation of response to osmotic stress), ion transporter activity, and ion homeostasis (Figure 4A).

Notably, LA significantly upregulated the expression of *CYCLIN D3-3* (Solyc04g078470) (Log2FC = 1.46), *CYCLIN D4-1* (Solyc11g010460) (Log2FC = 0.47), *NRT1.8* (Solyc07g008520) (Log2FC = 1.14), two *LYSINE HISTIDINE TRANSPORTER-LIKE 8* (*LHT8*, Solyc10g055740, Solyc04g082220) (Log2FC = 2.62; Log2FC = 1.32), *AMINO ACID TRANSPORTER* (Solyc05g052300) (Log2FC = 0.38), *CATIONIC AMINO ACID TRANSPORTER* (Solyc02g070280) (Log2FC = 0.57), *EXPANSIN A15 (EXPA15)* (Solyc05g052250) (Log2FC = 0.70), *HEAT SHOCK PROTEIN* 70 (HSP70) (Solyc04g011440) (Log2FC = 1.73), *HEAT SHOCK PROTEIN* 90 *(HSP90)* (Solyc06g036290) (Log2FC = 1.75), *HEAT SHOCK CHAPERON* (*HSC1,* Solyc06g076020) (Log2FC = 0.72), *HEAT SHOCK TRANSCRIPTION FACTOR* (Solyc09g065660) *(HSFA7)* (Log2FC = 1.16) (Table S7).

These genes are of particular interest: CYCLINS regulate the cell cycle progression (Potuschak and Doerner, 2001), *EXPA15* promotes cell expansion (Sampedro and Cosgrove, 2005; Lu *et al*., 2016), *HSFA7* responds at the early stages of HS (Mesihovic *et al*., 2022), *LHT8* encodes an AA transporter (Tegeder and Ward, 2012), and NRT1.8 functions as transporter of nitrate in vascular tissue (Li *et al*., 2010a). Additional *NRT1* family genes were also differentially expressed with *NRT1.1* (Solyc05g005990) (Log2FC = 0.32) upregulated and *NRT1.7* (Solyc08g007060) (Log2FC = -0.50), *NRT1.4* (Solyc04g005790) (Log2FC = -0.66) downregulated by LA (Table S7).

Then, DEGs specifically regulated by LA under HS were analyzed (Table S9). For the 1,131 downregulated genes, 115 GO terms were significantly enriched (Figure 4B, Table S10). These terms were associated with specialized cell wall development (suberin and lignin metabolic process), response to SA, biotic and abiotic stresses (response to fungus, response to wounding and salt, response to misfolded protein and antioxidant activity), transporter activity, and metabolism (specialized metabolism process, response to sucrose, fatty-acid derivate biosynthetic process)(Figure 4B). Downregulated genes belonging to the GO term antioxidant activity, included ten *PEROXIDASES* (Solyc01g067850, Solyc02g064970, Solyc02g094180, Solyc03g006700, Solyc05g010330, Solyc05g052280, Solyc06g050440, Solyc06g076630, Solyc11g018772, Solyc01g105070), and one *CATALASE* (Solyc12g094620) (Table S9). Also, LA-treated seedlings showed downregulation of genes belonging to GO lipid transport, including *NON-SPECIFIC LIPID TRANSFER PROTEIN* (Solyc01g081600, Solyc08g078030, Solyc09g065410, Solyc09g150148, Solyc09g082280), *LIPID TRANSFER PROTEIN* (Solyc06g054060, Solyc09g065420), Bidirectional sugar transporter *SWEET* (Solyc06g072620), *PHOSPHATIDYLGLYCEROL/ PHOSPHATIDYLINOSITOL TRANSFER PROTEIN* (Solyc10g081810). Independently of GO analyses, *ELONGATION OF FATTY ACIDS PROTEIN 3-LIKE* (Solyc03g044910) (Log2FC = -0.44) and *PROTEIN PHOSPHATASE 2C* (Solyc03g096670) (Log2FC = -1.37) and *FATTY ACID DESATURASE* (Solyc04g040120) (Log2FC = -0.85) were downregulated (Table S9). For the 677 upregulated genes, 18 GO BP terms were significantly enriched (Figure 4B, Table S10). These were primarily related to specific metabolic processes (xyloglucan metabolism process, polysaccharide catabolic process), cellular response to nutrient levels, regulation of systemic acquired resistance, and detoxification.

Overall, the RNA-Seq analysis revealed that LA stimulated the expression of cell division markers under both NS and HS conditions, potentially explaining the observed increase in the division zone length of the primary root apical meristem. The upregulation of *NRT1.8* under NS, together with the enrichment of GO terms related to nutrient levels, hormonal signaling, and different metabolic pathways, suggests that LA may also be involved in modifying sugar and lipid metabolism of seedlings. Interestingly, *HSFA7* and *HSC70* were upregulated by LA under NS, supporting that the treatment may prime seedlings for abiotic stress (Figure 4, Table S7).

### LA modulates phytohormones associated with root growth and abiotic stress responses

The transcriptomic data suggest an impact of LA on the hormonal pathways under NS (Figure 4). Phytohormones such as auxin, ABA, CKs, brassinosteroid (BRs), JA, GA, and SA are key regulators of root growth *via* their action on cell division and/or expansion (Del Pozo *et al*., 2005; Li *et al*., 2015; Li, Sun and Liu, 2022). HS also alters hormone homeostasis and spatial distribution (Tiwari *et al*., 2022). To determine whether LA affects the hormonal status of seedlings, the levels of seven hormones (ABA, GA, JA, jasmonate-isoleucine (JA-Ile), MeJA, SA, and CK) were quantified in roots collected 12 days after sowing, corresponding to 5 days after transfer to LA medium under NS.

LA significantly increased JA, JA-ILe, and SA levels by 57%, 74%, and 56%, respectively. Conversely, MeJA level significantly decreased by 76% (Figure 5). In contrast, LA did not significantly affect ABA, Zeatin, nor GA levels (Figure 5).

**Figure 5.**
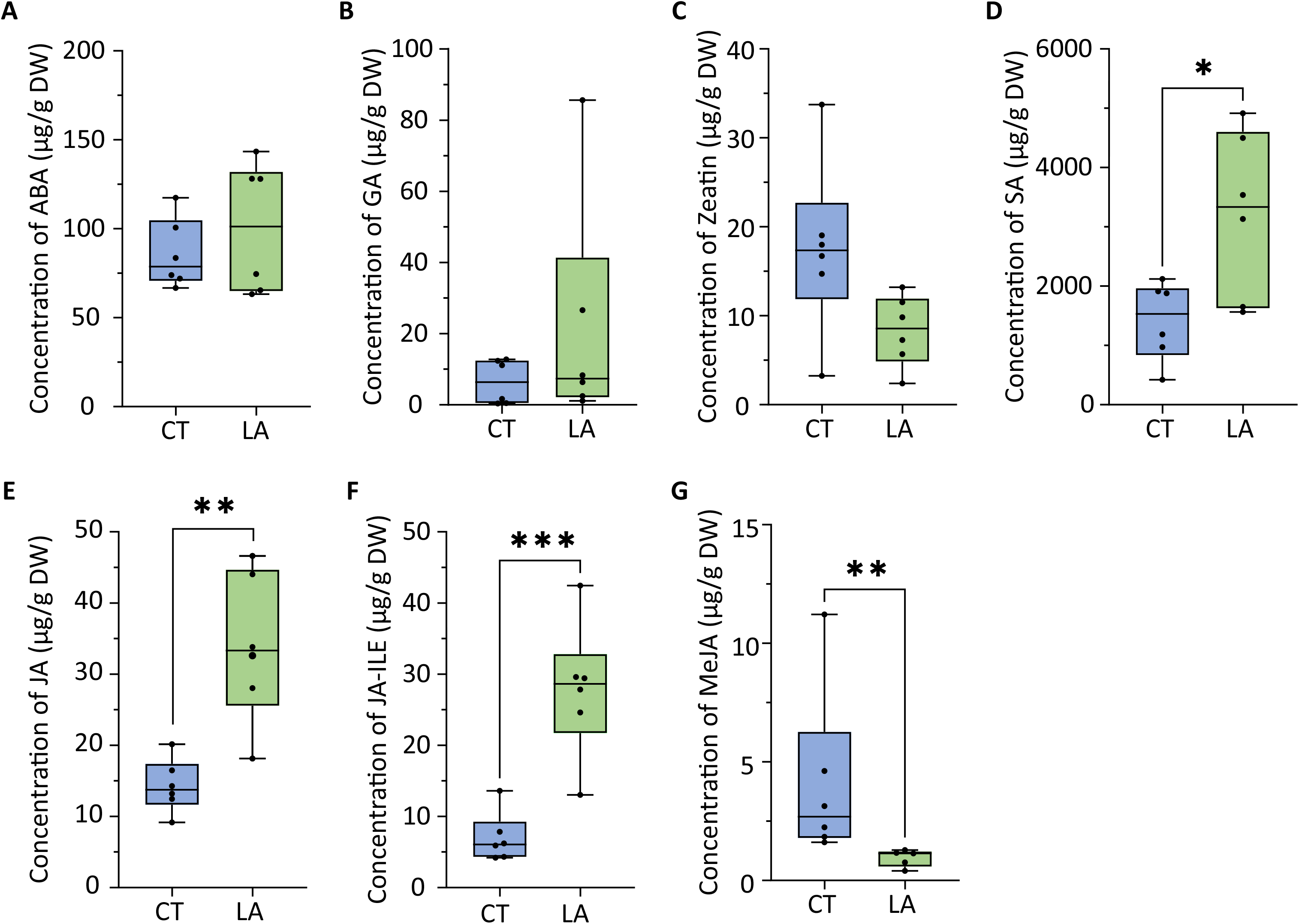
LA treatment under NS conditions modifies the levels of phytohormones related to plant defense against biotic and abiotic stresses. Quantified hormones include (A) abscisic acid (ABA), (B) gibberellic acid (GA), (C) zeatin, (D) salicylic acid (SA), (E) jasmonic acid (JA), (F) jasmonoyl-isoleucine (JA-Ile), and (G) methyl jasmonate (MeJA). Boxplot: whiskers extend from minimum to maximum, box extends 25th to 75th percentiles, the line in the middle is the median. Asterisks indicate statistically significant differences between treated and untreated plants. Statistical analysis was performed using a t-test; *p < 0.05, **p < 0.005, ***p < 0.0005, and ****p < 0.0001.

The transcriptomic data were then interrogated to investigate whether alterations in hormone levels were reflected in the expression of hormone-related genes. First, we defined hormonal biosynthetic pathways based on Arabidopsis thaliana homologs and the KEGG database (Kanehisa and Goto, 2000; Peng *et al*., 2021) (Figure S5) and then identified homologous genes in our transcriptomic dataset. Several genes involved in SA, JA, JA-Ile, and MeJA biosynthetic pathways were differentially expressed in response to LA (Figure S5). In the SA pathway, two genes encoding for SA biosynthesis were upregulated: *PHENYLALANINE AMMONIA-LYASE (PAL)* (Solyc00g500095), and *ABNORMAL INFLORESCENCE MERISTEM 1 (AIM1)* (Solyc07g019670); and one gene was downregulated: *ENHANCED PSEUDOMONAS SUSCEPTIBILITY 1 (EPS1)* (Solyc01g107070). Regulatory genes of the SA signaling pathway were also modulated, such as *SYSTEMIC ACQUIRED RESISTANCE DEFICIENT 1 (SARD1)* (Solyc03g119250) and *WRKY DNA-binding protein 46 (WRKY46)* (Solyc01g095630), *NPR1-Interacting Protein 1 (NIMIN1)* (Solyc01g100940), *NONEXPRESSOR OF PATHOGENESIS-RELATED GENES 3 (NRP3)* (Solyc02g069310), and *WRKY DNA-BINDING PROTEIN 70 (WRKY70)* (Solyc03g095770) which were upregulated, and *ANAC019* (Solyc07g063410), *GLUTAMATE SYNTHASE 1 (GLT1)* (Solyc03g083440) which were downregulated (Figure S5A) (Peng *et al*., 2021). In the JA biosynthetic pathway, only two genes, *AIM1* (Solyc07g019670) and *ACETYL-COA ACYLTRANSFERASE* (Solyc02g064690), were upregulated (Figure S5B).

Overall, these results indicated that the LA modulated stress-related hormonal pathways in roots under NS, by promoting the accumulation of JA, JA-Ile, and SA, while reducing MeJA levels, known to inhibit primary root growth. These results support the hypothesis that LA could prime seedlings for abiotic stress.

### LA induces widespread metabolic reprogramming, stimulating sugar, fatty acid, and the ornithine-citrulline pathway

Similarly, transcriptomic data suggest an impact of LA on different metabolic pathways under NS (Figure 4). To explore the impact of LA on metabolism, roots were collected 12 days after sowing, corresponding to 5 days after transfer to LA medium under NS, and metabolic profiles of LA and CT seedlings were compared by GC-MS. Levels of confidence in metabolite identification is described in Table S11.

A total of 55 metabolites were identified and quantified with a polar extraction (Table S12). PCA of the identified polar metabolites revealed a strong LA effect, with treated samples forming a distinct cluster, accounting for 39% of the variance (Figure 6A). LA-treated roots showed increased levels of non-amino acid nitrogen-containing compounds (cadaverine, pyroglutamic acid) (LA/CT = 2.08; LA/CT = 6.16), organic acids (galactaric acid, α-ketoglutaric acid (AKG), lactic acid) (LA/CT = 1.28; LA/CT = 12.45; LA/CT = 2.68), and sugars (fructofuranose, fructofuranoside, galactose, hexose, mannose, ribose, sucrose, xylose) (LA/CT = 4.38; LA/CT > 10; LA/CT = 0.61; LA/CT = 32.18; LA/CT = 2.94; LA/CT = 6.85; LA/CT = 1.64; LA/CT = 3.99; LA/CT = 7.61). Most AA levels were unaffected by LA, except for increases in valine, isoleucine, citrulline, and γ-aminobutyric acid (GABA), and a decrease in proline (Table S13). From the non-polar extract, 54 metabolites were identified (Table S12). PCA of the non-polar metabolites did not clearly separate LA-treated from CT root (Figure 6B). However, some metabolites displayed significant increased abundance following treatment (LA/CT > 1), including: fatty acid derivatives (C16:0-glycerol, C18:0-glycerol, C17:0, hexadecanoic acid), non-amino acid N-containing compound (putrescine), polyol (glycerol), carboxylic acid (phenylpropanoid), and organic acid (benzoic acid) (Table S13).

**Figure 6.**
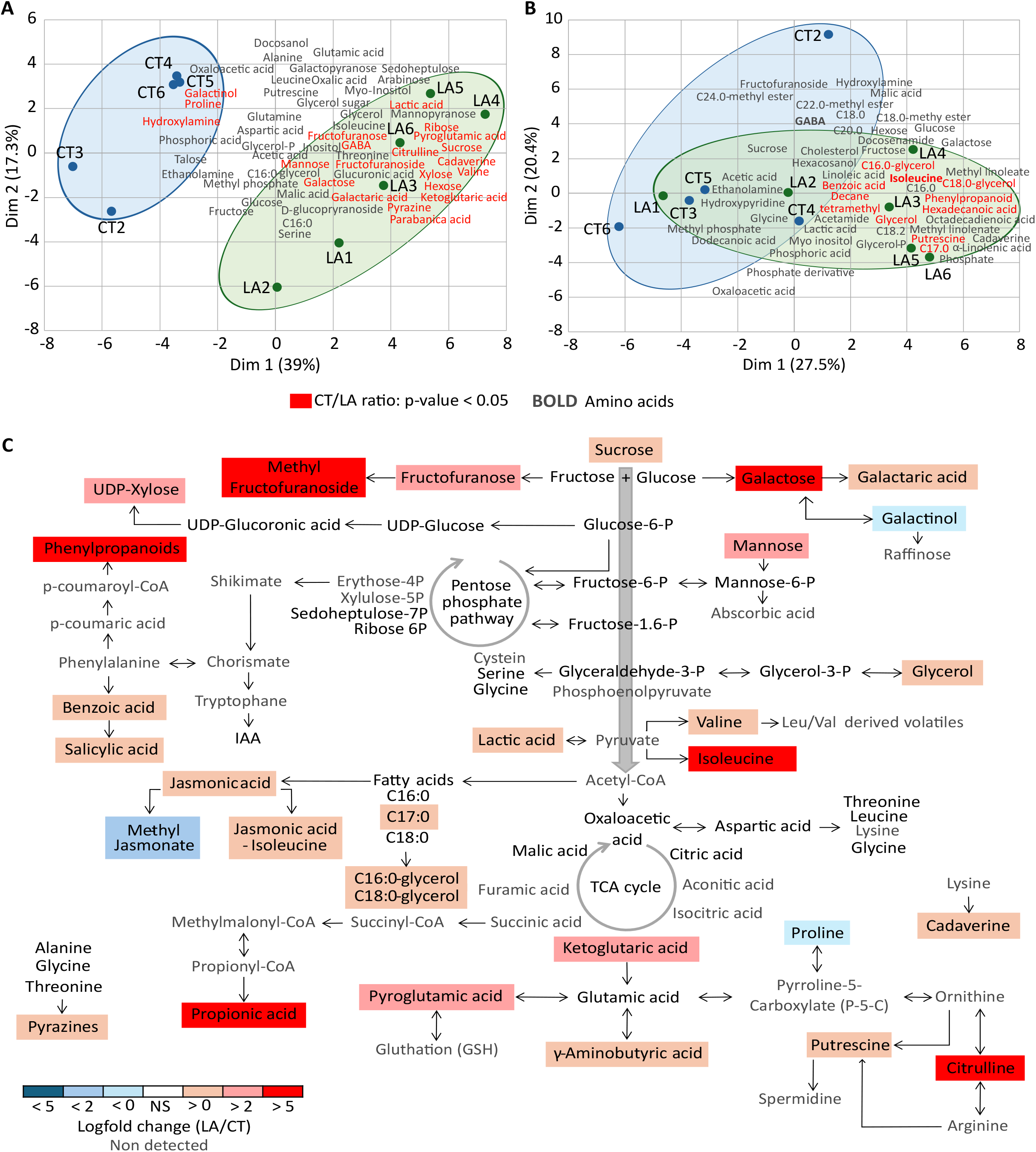
LA treatment under NS conditions induces metabolic reprogramming of tomato roots. (A) Biplot of metabolic changes triggered by the LA treatment. PCA using all detected metabolites obtained by polar extraction. (B) Biplot of metabolic changes triggered by the LA treatment. PCA using all detected metabolites obtained by non-polar extraction. Ratio and exact p-values for each metabolite are provided in Table S12 and Table S13. Amino acids are in bold, and compounds significantly influenced by the LA treatment are labelled in red, p < 0.05. (C) Biochemical pathways summarised metabolite changes triggered by the LA treatment. Metabolite levels are shown relative to untreated roots and expressed as log₂ fold change (LA/CT). Colours indicate statistically significant changes, which are highlighted using the corresponding colour and intensity of the fold change. Detected metabolites are shown in black, and undetected ones in grey. Amino acids are indicated in bold.

To obtain a global perception on the metabolic adjustments, metabolite and phytohormone ratios were mapped onto the central metabolic pathways (Figure 6C). LA-treated roots exhibited widespread increased metabolite levels across primary metabolic pathways. Notable trends included: higher sugar levels, minimal impact on the tricarboxylic acid (TCA) cycle, and increased levels of 2 metabolites in the ornithine-citrulline pathway. Taking into account the phytohormone profiling (Figure 5), it seems that the LA triggered an upregulation of JA biosynthesis, supported by enhanced fatty acid metabolism required for its synthesis; an increased SA biosynthesis despite stable levels of some intermediates; and a similar pattern in the phenylpropanoid pathway, where the end products of the pathway were up-regulated despite no difference in intermediary metabolite levels being observed.

Together, the metabolomic and hormonal data reveal that LA induced coordinated metabolic reprogramming in roots, enhancing sugar metabolism and stress-related phytohormone pathways. These changes, combined with the upregulation of HS response markers under NS, suggest a priming effect of LA.

### LA might prepare the plants to withstand HS

To determine whether LA induces a priming effect, an experiment in which the treatment was applied before exposing plants to HS was designed. Three-day-old tomato seedlings were transferred onto either MS medium or MS supplemented with LA. After five days, all plants grown on MS medium were transferred to fresh MS medium (MS), while LA-treated seedlings were divided into two groups: one transferred to fresh MS medium (LA P, for “primed”), and the other transferred to fresh MS medium containing LA (LA). Immediately after transfer, all plants were subjected to HS, and primary root length was recorded daily for seven days (Figure 7A). Plants continuously treated with LA before and during the HS (LA) showed significantly longer primary roots (27% longer at the end of the experiment) compared with MS seedlings. Notably, LA P seedlings also displayed increased primary root length (18% by the end of the experiment) relative to MS seedlings (Figure 7B) (Table S14). These results demonstrate that LA pre-treatment alleviated heat induced inhibition of root growth, supporting the hypothesis that LA primes seedlings to better withstand subsequent HS.

**Figure 7.**
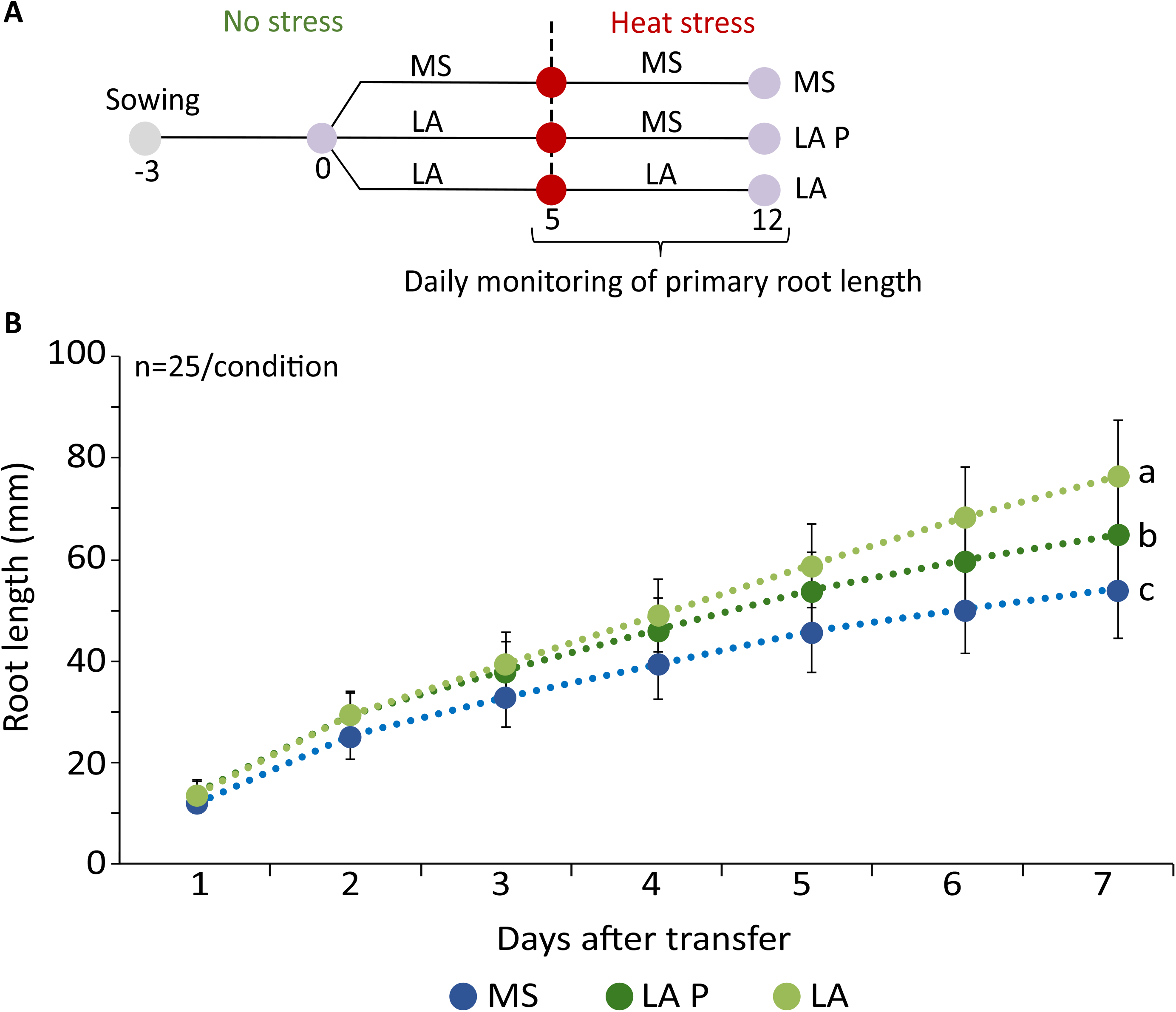
Evaluation of the priming effect of LA treatment and its impact on plant tolerance to HS. (A) Experimental design used to assess the priming effect of LA treatment prior to heat stress. (B) Primary root length of tomato seedlings over time under HS, following different timings of LA application. Data represent mean ± SD. Letters indicate statistically significant differences compared to untreated plants (p < 0.05). Exact p-values for each comparison at each time point are provided in Table S12.

## Discussion

### LA promotes root growth by stimulating cell division, potentially contributing to cell expansion, and influencing nutrient allocation

Primary root development relies on a finely tuned balance between cell production through cell division and subsequent cell expansion, two processes that are spatially separated by the transition zone (Dello Ioio *et al*., 2007; Desvoyes, Echevarría and Gutierrez, 2021). Previous studies have shown that exogenous AA can increase root meristem size, with compounds such as proline, asparagine, glutamate, and methionine (Biancucci *et al*., 2015). LA appeared to act in a similar manner since its application enhanced root growth by increasing root diameter, the length of the division zone, and potentially the length of the elongation zone, suggesting an effect on both cell division and expansion (Figure 2).

At the molecular level, LA upregulated the expression of D-type cyclins, particularly *CYCD3;3* (Table S7), which plays a pivotal role in early root development by mediating G1/S cell cycle transition and promoting localized cell proliferation (Riou-Khamlichi *et al*., 1999; Dewitte *et al*., 2007). Together with other cyclins, *CYCD3;3* is essential for sustaining cell proliferation and for the correct establishment of the QC (Forzani *et al*., 2014). *EXPANSIN* genes are known to be involved in cell expansion and cell-wall modification (Li, Jones and McQueen-Mason, 2003). In Arabidopsis, *EXPA15* is localized in the epidermis of RAM and in the emerging lateral root. *EXPA15* is identified as an upregulated gene in the gain-of-function *edt1* mutant, and is associated with the development of deeper primary roots (Xu *et al*., 2014; Samalova *et al*., 2020). LA induced the expression of the cell division marker *CYCD3;3* (Log2FC = 1.46) and the cell expansion marker *EXPA15* (Log2FC = 0.70), suggesting that LA modulated cell division and expansion through transcriptional regulation. In the primary root, nitrate can promote growth by enhancing meristematic activity (Naulin *et al*., 2020). The nitrate transporter family is a large group of transporters that are well characterized for their role in nitrate uptake. NRT1.1 is a dual-affinity nitrate transporter that not only mediates both high-and low-affinity uptake but also senses the amount of nitrate available (Ho *et al*., 2009). The increased expression of *NTR1.1* and *NRT1.8* (Log2FC = 0.32 or 1.14, respectively), which retrieve nitrate from xylem vessels, suggests that LA could modulate nitrate transport (Li *et al*., 2010b; Sun *et al*., 2015). The downregulation of *NRT1.7* (Log2FC = -0.50), involved in phloem loading, and *NRT1.4* (Log2FC = -0.66), the plasma membrane nitrate transporter, further supports this hypothesis (Fan *et al*., 2009; Morales de los Ríos *et al*., 2021). Similarly, *LHT* transporters, found upregulated by LA, are known to be essential for intracellular recycling and import of organic nitrogen Tegeder and Ward, (2012); Sestili *et al*., (2018) have already demonstrated that under high nitrogen levels, the application of a PHs increased nitrate concentration in tomato leaves. The increase of organic acid and expression of several nitrate, sulfur, and phosphate transporters, *OPT9* in LA-treated roots (Table S7), suggests that LA may enhance nutrient redistribution, including modulation of organic nitrogen allocation, which can impact root growth.

### LA mitigates the HS effect on roots

HS triggers oxidative stress in plants, leading to the excessive accumulation of ROS (O₂⁻, H₂O₂), causing cellular damage, while H₂O₂ also acts as a secondary messenger mediating stress signaling. To counteract ROS toxicity, plants deploy enzymatic antioxidants (CAT, POD, SOD) and non-enzymatic compounds (ascorbate, glutathione, phenolics) (Gill and Tuteja, 2010). LA-treated roots under HS showed downregulation of antioxidant activity-related genes, including ten POD and one CAT compared to untreated HS seedling (Table S8, Figure 4B), supporting the notion that LA limits ROS accumulation under HS, thereby attenuating the need for antioxidant gene upregulation and suggesting an upstream effect by reducing oxidative pressure rather than enhancing enzymatic detoxification systems. This is consistent with the results from Ertani *et al*., (2013), who reported decreased antioxidant enzyme activities following PH application under salinity in maize, though in contrast, AA mixtures have been shown to increase POD and CAT activities in soybean (Teixeira *et al*., 2017). HS alters membrane lipid composition by inducing changes in lipid saturation, fatty acid chain length (Niu and Xiang, 2018), and cell wall integrity by compromising mechanical stability (Wu, Bulgakov and Jinn, 2018). Under HS, LA-treated seedlings showed downregulation of genes involved in lipid transport and fatty acid metabolism (*NON-SPECIFIC LIPID-TRANSFER PROTEIN, 2 LIPID TRANSFER PROTEIN, ELONGATION OF FATTY ACIDS PROTEIN 3-LIKE, PROTEIN PHOSPHATASE 2C*, *FATTY ACID DESATURASE)*, suggesting reduced heat induced membrane damage and a lower need for membrane remodeling (Table S9, S10). Regarding the cell wall, LA triggered the upregulation of 21 extensins while wax, suberin, lignin, and cutin biosynthesis genes were downregulated (Figure 4B, Table S10). This pattern indicates a reallocation in cell wall construction from hydrophobic barriers (wax, suberin, cutin, lignin) involved in rigidification toward a more dynamic, extensin-mediated reinforcement. As structural proteins involved in wall rigidification through ROS-dependent crosslinking (Moussu and Ingram, 2023), extensin upregulation suggests that LA reinforced cell wall integrity while maintaining plasticity. Together with the upregulation of *CYCD3;3* (Log2FC = 0.94) and *EXPA15* (Log2FC = 0.49) (Table S9), and enhanced primary metabolism, this cell wall remodeling is consistent with sustained cell division and expansion under HS.

### LA prepares the plant to withstand HS

Priming is a physiological state enabling plants to respond faster and more effectively to subsequent biotic or abiotic stresses (Conrath *et al*., 2006; Aranega-Bou *et al*., 2014). Triggered by diverse stimuli, including abiotic factors and chemical compounds, priming induces physiological, molecular, and metabolic changes that prepare plants for future stress, not necessarily of the same nature as the initial stimulus (Hilker *et al*., 2016; Mauch-Mani *et al*., 2017).

Seedlings pre-treated with LA exhibited sustained growth with delayed impairment under HS, suggesting a favourable trade-off between growth and stress resilience (Figure 7B) (Conrath *et al*., 2006). Consistently, transcriptomic analyses revealed enrichment of protein folding and HS response GO categories in LA-treated root under NS, with upregulation of HS markers, indicating that protective mechanisms were activated before stress exposure, as illustrated by the enriched GO terms related to stress response (Figure 4, Table S7).

The upregulation of *HSFA7*, which initially acts as a co-repressor and later as a co-activator of *HSFA1a*, to fine-tune the HS response (Mesihovic *et al*., 2022), and the upregulation of *HSP70* and *HSP90*, involved in proteasomal degradation of unfolded proteins and stabilization, respectively (Pratt *et al*., 2010; Hahn *et al*., 2011), by LA treatment under NS, suggested that LA contributes to long-term resilience against HS. *HSP16.9, HSP18.1, HSP22,* and *HSP26* were also upregulated by a PH in maize roots under NS. The authors suggested that this response might reflect a priming effect of the biostimulant (Vaseva *et al*., 2022).

LA increased JA and SA concentrations in roots under NS (Figure 5). This finding supports the priming effect of LA since it is consistent with a study describing that JA application can mitigate drought stress effects, stimulate antioxidant activity in sunflowers. Seed priming is also induced by exogenous application of SA and JA under HS (Górnik, Badowiec and Weidner, 2014; Ashraf and Siddiqi, 2024). Additionally, SA may modulate genes involved in the cell cycle and cell expansion, influencing root meristem activity and elongation. The overall effect of SA on root growth is dependent on the plant species, organ type, and physiological context (Pasternak *et al*., 2019; Bagautdinova *et al*., 2022; Li, Sun and Liu, 2022). In our study, an increase of SA concentration in LA-treated roots correlated with enhanced root growth, consistent with the tomato-specific responses to HS reported by Khan *et al*., (2019) (Figure 5D). LA also upregulated PAL, a key enzyme in SA biosynthesis, along with SA-responsive genes WRKY46 and WRKY70 (Hu, Dong and Yu, 2012) (Table S7, Figure S5), consistent with PH and AA studies in maize and soybean (Ertani *et al*., 2013; Teixeira *et al*., 2017; Canellas *et al*., 2025). Although JA typically inhibits proximal meristem cell division in Arabidopsis (Chen *et al*., 2011). The accumulation of JA, MeJA, and JA-Ile in LA-treated roots without growth impairment further supports a priming effect, illustrating how context-dependent hormonal signaling can enhance stress defense without compromising development. This hormonal reprogramming may contribute to the LA-induced enhancement of root development and stress tolerance.

Metabolomic profiling further supports a priming effect of LA. LA elevated citrulline (LA/CT > 10), associated with drought tolerance (Kawasaki *et al*., 2000; Song *et al*., 2020), and putrescine (LA/CT = 1.73), a polyamine enhancing antioxidant activity under drought (Zhao *et al*., 2021; Islam *et al*., 2022). Additionally, GABA, AKG, phenylpropanoids, and pyroglutamic acid were also increased: GABA provides osmoprotection and ROS scavenging under multiple abiotic stresses (Abd Elbar *et al*., 2021; Abdel Razik *et al*., 2021; Hasan *et al*., 2021), while AKG enhances heat tolerance through antioxidant reinforcement and membrane stability (Lei and Huang, 2022). In addition, LA induced accumulation of phenylpropanoids, described to be accumulated and to serve as ROS scavengers under temperature stress (Ninkuu *et al*., 2025), consistent with previously described PH modes of action (Colla *et al*., 2015). Their upregulation by LA treatment under NS may therefore act as a priming mechanism, preparing seedlings to better cope with subsequent abiotic stresses (Figure 6, Table S13, 14).

Moreover, under HS, membrane rigidity is maintained through increased proportions of saturated and di-unsaturated fatty acids (Falcone, Ogas and Somerville, 2004). The LA-induced accumulation of palmitic and stearic acids under NS may therefore prepare the cell membrane to adapt its fluidity in function of environmental conditions, further supporting a LA-induced priming effect.

Collectively, these results demonstrate that Leafamine® enhances tomato root resilience to heat stress through a dual mode of action; direct mitigation of heat induced cellular damage and the establishment of a primed physiological state that prepares seedlings for future stress. The underlying cellular processes contributing to these mode of actions include; the limitation of ROS accumulation, which reinforces membrane integrity and cell wall stability, while modulating key hormonal and metabolic pathways, LA reduces the cellular cost of stress adaptation and preserves root growth under high temperatures. The activation of HS-related genes, accumulation of protective metabolites, and hormonal reprogramming under NS further support the existence of a robust priming effect. These findings not only clarify the molecular and metabolic mechanisms underlying LA activity but also position protein hydrolysate-based biostimulants as promising tools to enhance crop resilience in a warming climate. Heat stress is becoming an increasingly critical constraint on agricultural productivity, LA may offer a sustainable and effective strategy to safeguard root function and support plant performance under future environmental conditions.

## Supplemental Data

**Supplemental Figure 1.** Dose response of LA treatment on primary root development of tomato seedlings.

**Supplemental Figure 2.** Effect of the LA treatment on shoot growth of tomato seed-lings grown under NS and heat stress HS conditions for 7 days.

**Supplemental Figure 3.** Effect of LA on the length of the elongation zone of the primary root of seedlings grown under NS and HS conditions.

**Supplemental Figure 4.** LA treatment exerts condition-dependent effects under HS, specifically impacting molecular processes related to metabolism and stress responses.

**Supplemental Figure 5.** LA treatment exerts condition-dependent effects under HS, specifically impacting molecular processes related to metabolism and stress response.

**Supplemental Table 1.** Results of the ANOVA test performed to evaluate the effects of different concentrations of LA on primary root growth.

**Supplemental Table 2.** Results of the ANOVA test performed to evaluate the effects of heat stress and LA on primary root growth.

**Supplemental Table 3.** List of DEGs resulting from the LA x HS interaction, clustered according to expression profile.

**Supplemental Table 4.** GO enrichment analysis of each cluster resulting from the LA x HS interaction.

**Supplemental Table 5.** List of common up-and down-regulated genes by LA treatment under NS and HS conditions.

**Supplemental Table 6.** GO enrichment analysis of common up-and down-regulated genes by LA treatment under NS and HS conditions.

**Supplemental Table 7.** List of DEGs induced by LA treatment under NS conditions.

**Supplemental Table 8.** GO enrichment analysis of DEGs induced by LA treatment under NS conditions.

**Supplemental Table 9.** List of DEGs induced by LA treatment under HS conditions.

**Supplemental Table 10.** GO enrichment analysis of DEGs induced by LA treatment under HS conditions.

**Supplemental Table 11.** Confidence levels of metabolite identification according to MSI.

**Supplemental Table 12.** Polar metabolite profiles of tomato roots treated with LA under NS conditions.

**Supplemental Table 13.** Non-polar metabolite profiles of tomato roots treated with LA under HS conditions.

**Supplemental Table 14.** Results of the ANOVA test performed on priming effects of LA treatments on primary root growth.

## Acknowledgements

This work was carried out with the financial support of ANRT (National Association for Research and Technology) (n°2022/1477). The microscopy analyses were performed in the Bordeaux Imaging Center, a service unit of the CNRS-INSERM and Bordeaux University, member of the national infrastructure France BioImaging. We express our deepest thanks to Isabelle Atienza for taking care of the plant culture in the greenhouse.

## Author contribution

L.M; E.M; M.H conceived the project and designed the research. L.M; and N.B performed the research. L.M; M.N; P.D.F; analyzed the hormonal and metabolite results. All authors analyzed and discussed the results. All authors wrote the manuscript with input from the other authors.

## Data availability statement

The RNA-seq data have been deposited in ArrayExpress under accession number E-MTAB-17336.

## Figure legends

**Figure S1.** Dose response of LA treatment on primary root development of tomato seedlings. (A) Schematic representation of a tomato plantlet and the time points for root length measurement. (B) Experimental design of the dose-response assay for LA treatment. (C) Primary root growth over time in seedlings treated with different LA concentrations. (D) Representative images of plates with seedlings grown on media supplemented with increasing concentrations of LA. Scale bar = 8 cm. Statistical comparisons (p-value) are shown in Table S2. Letters indicate significant differences between treatments. ANOVA test was used followed by Tukey’s multiple comparisons test.

**Figure S2.** Effect of the LA treatment on shoot growth of tomato seedlings grown under NS and heat stress HS conditions for 7 days. (A) Shoot fresh weight. (B) Shoot dry weight. (C) Shoot fresh-to-dry weight ratio (FW/DW). Boxplot: whiskers extend from minimum to maximum, box extends 25th to 75th percentiles, the line in the middle is the median. Letters indicate significant differences among treatments. Asterisks indicate significant differences. ANOVA test was used followed by Tukey’s multiple comparisons test; *p < 0.05, p < 0.005, *p < 0.0005, ****p < 0.0001.

**Figure S3.** Effect of LA on the length of the elongation zone of the primary root of seedlings grown under NS and HS conditions. (A, C, E, G) Primary root tip of untreated, unstressed plants (CT) (A); LA-treated, unstressed plants (LA) (C); untreated, heat-stressed plants (HS) (E); LA-treated, heat stressed plant (LAHS) (G). Scale bar = 2mm. (B, D, F, H) Close-up view (white rectangle in A, C, E, G) of the region presenting the upper boundary of the elongation zone for each condition, defined as the position of the first root hair (white Arrows). Scale bar = 200µm

**Figure S4.** LA treatment exerts condition-dependent effects under HS, specifically impacting molecular processes related to metabolism and stress responses. (A) GO enrichment analysis of each cluster resulting from the LA x HS interaction. (B) GO enrichment analysis of DEGs (Log2FC > 0.5) consistently up-or downregulated by LA treatment, under both NS and HS conditions. Dvl = Development. Hor = Hormones. Metabo = Metabolism. Nu = Nutrition. Ot = Others. Trans = Transcription. GO terms are grouped by category; dot color represents adjusted p-values, and dot size indicates gene ratio. GO terms from table S4, table S5, and table S6 were grouped to remove redundancy.

**Figure S5.** Schemes representing the transcriptional changes of phytohormone biosynthesis pathways in roots five days after LA treatment under NS conditions based on the transcriptomic data. (A) Expression changes in genes involved in the SA biosynthetic pathway. (B) Expression changes in genes involved in the JA biosynthetic pathway. The schematic shows key enzymes and intermediates of each pathway. Colour shading represents the log₂ fold-change ratio (CT/LA) in gene expression, with red indicating upregulation and blue indicating downregulation. Genes with statistically significant expression changes (p < 0.05) are marked by coloured boxes. The colour scale bar indicates the magnitude and direction of the change.

## Reference

Abd Elbar, O.H., et al. (2021) ‘Protective Effect of γ-Aminobutyric Acid Against Chilling Stress During Reproductive Stage in Tomato Plants Through Modulation of Sugar Metabolism, Chloroplast Integrity, and Antioxidative Defense Systems’, Frontiers in Plant Science, 12, p. 663750. Available at: 10.3389/fpls.2021.663750.

Abdel Razik, E.S., et al. (2021) ‘γ-Aminobutyric acid (GABA) mitigates drought and heat stress in sunflower (Helianthus annuus L.) by regulating its physiological, biochemical and molecular pathways’, Physiologia Plantarum, 172(2), pp. 505–527. Available at: 10.1111/ppl.13216.

Alsamir, M., et al. (2021) ‘An overview of heat stress in tomato (Solanum lycopersicum L.)’, Saudi Journal of Biological Sciences, 28(3), pp. 1654–1663. Available at: 10.1016/j.sjbs.2020.11.088.

Aranega-Bou, P., et al. (2014) ‘Priming of plant resistance by natural compounds. Hexanoic acid as a model’, Frontiers in Plant Science, 5. Available at: 10.3389/fpls.2014.00488.

Ashraf, F. and Siddiqi, E.H. (2024) ‘Mitigation of drought-induced stress in sunflower (Helianthus annuus L.) via foliar application of Jasmonic acid through the augmentation of growth, physiological, and biochemical attributes’, BMC Plant Biology, 24(1), p. 592. Available at: 10.1186/s12870-024-05273-4.

Bagautdinova, Z.Z., et al. (2022) ‘Salicylic Acid in Root Growth and Development’, International Journal of Molecular Sciences, 23(4), p. 2228. Available at: 10.3390/ijms23042228.

Biancucci, M., et al. (2015) ‘Proline affects the size of the root meristematic zone in Arabidopsis’, BMC Plant Biology, 15(1), p. 263. Available at: 10.1186/s12870-015-0637-8.

Campobenedetto, C., et al. (2021) ‘A Biostimulant Based on Seaweed (Ascophyllum nodosum and Laminaria digitata) and Yeast Extracts Mitigates Water Stress Effects on Tomato (Solanum lycopersicum L.)’, Agriculture, 11(6), p. 557. Available at: 10.3390/agriculture11060557.

Canellas, L.P., et al. (2025) ‘Farm-Produced Plant Biostimulant: Case Study with Passion Fruit’, Agronomy, 15(3), p. 681. Available at: 10.3390/agronomy15030681.

Ceccarelli, A.V., et al. (2021) ‘Foliar Application of Different Vegetal-Derived Protein Hydrolysates Distinctively Modulates Tomato Root Development and Metabolism’, Plants, 10(2), p. 326. Available at: 10.3390/plants10020326.

Chen, Q., et al. (2011) ‘The Basic Helix-Loop-Helix Transcription Factor MYC2 Directly Represses PLETHORA Expression during Jasmonate-Mediated Modulation of the Root Stem Cell Niche in Arabidopsis’, The Plant Cell, 23(9), pp. 3335–3352. Available at: 10.1105/tpc.111.089870.

Colla, G. et al. (2014) ‘Biostimulant action of a plant-derived protein hydrolysate produced through enzymatic hydrolysis’, Frontiers in Plant Science, 5. Available at: https://www.frontiersin.org/articles/10.3389/fpls.2014.00448 (Accessed: 19 August 2022).

Colla, G., et al. (2015) ‘Protein hydrolysates as biostimulants in horticulture’, Scientia Horticulturae, 196, pp. 28–38. Available at: 10.1016/j.scienta.2015.08.037.

Conrath, U., et al. (2006) ‘Priming: Getting Ready for Battle’, Molecular Plant-Microbe Interactions®, 19(10), pp. 1062–1071. Available at: 10.1094/MPMI-19-1062.

Del Pozo, J.C., et al. (2005) ‘Hormonal control of the plant cell cycle’, Physiologia Plantarum, 123(2), pp. 173–183. Available at: 10.1111/j.1399-3054.2004.00420.x.

Dello Ioio, R., et al. (2007) ‘Cytokinins Determine Arabidopsis Root-Meristem Size by Controlling Cell Differentiation’, Current Biology, 17(8), pp. 678–682. Available at: 10.1016/j.cub.2007.02.047.

Desvoyes, B., Echevarría, C. and Gutierrez, C. (2021) ‘A perspective on cell proliferation kinetics in the root apical meristem’, Journal of Experimental Botany, 72(19), pp. 6708– 6715. Available at: 10.1093/jxb/erab303.

Dewitte, W., et al. (2007) ‘Arabidopsis CYCD3 D-type cyclins link cell proliferation and endocycles and are rate-limiting for cytokinin responses’, Proceedings of the National Academy of Sciences, 104(36), pp. 14537–14542. Available at: 10.1073/pnas.0704166104.

Dobin, A., et al. (2013) ‘STAR: ultrafast universal RNA-seq aligner’, Bioinformatics, 29(1), pp. 15–21. Available at: 10.1093/bioinformatics/bts635.

Ertani, A., et al. (2013) ‘Alfalfa plant-derived biostimulant stimulate short-term growth of salt stressed Zea mays L. plants’, Plant and Soil, 364(1–2), pp. 145–158. Available at: 10.1007/s11104-012-1335-z.

Falcone, D.L., Ogas, J.P. and Somerville, C.R. (2004) ‘Regulation of membrane fatty acid composition by temperature in mutants of Arabidopsis with alterations in membrane lipid composition’, BMC Plant Biology, 4, p. 17. Available at: 10.1186/1471-2229-4-17.

Fan, S.-C., et al. (2009) ‘The Arabidopsis Nitrate Transporter NRT1.7, Expressed in Phloem, Is Responsible for Source-to-Sink Remobilization of Nitrate’, The Plant Cell, 21(9), pp. 2750–2761. Available at: 10.1105/tpc.109.067603.

Forzani, C., et al. (2014) ‘WOX5 Suppresses CYCLIN D Activity to Establish Quiescence at the Center of the Root Stem Cell Niche’, Current Biology, 24(16), pp. 1939–1944. Available at: 10.1016/j.cub.2014.07.019.

Gill, S.S. and Tuteja, N. (2010) ‘Reactive oxygen species and antioxidant machinery in abiotic stress tolerance in crop plants’, Plant Physiology and Biochemistry, 48(12), pp. 909–930. Available at: 10.1016/j.plaphy.2010.08.016.

Giri, A., et al. (2017) ‘Heat Stress Decreases Levels of Nutrient-Uptake and -Assimilation Proteins in Tomato Roots’, Plants, 6(1), p. 6. Available at: 10.3390/plants6010006.

González-García, M.P., et al. (2023) ‘Temperature changes in the root ecosystem affect plant functionality’, Plant Communications, 4(3), p. 100514. Available at: 10.1016/j.xplc.2022.100514.

Górnik, K., Badowiec, A. and Weidner, S. (2014) ‘The effect of seed conditioning, short-term heat shock and salicylic, jasmonic acid or brasinolide on sunflower (Helianthus annuus L.) chilling resistance and polysome formation’, Acta Physiologiae Plantarum, 36(10), pp. 2547–2554. Available at: 10.1007/s11738-014-1626-5.

Guo, T., et al. (2022) ‘Heat stress mitigation in tomato (Solanum lycopersicum L.) through foliar application of gibberellic acid’, Scientific Reports, 12(1), p. 11324. Available at: 10.1038/s41598-022-15590-z.

Hahn, A., et al. (2011) ‘Crosstalk between Hsp90 and Hsp70 Chaperones and Heat Stress Transcription Factors in Tomato’, The Plant Cell, 23(2), pp. 741–755. Available at: 10.1105/tpc.110.076018.

Haider, S., et al. (2022) ‘Analyzing the regulatory role of heat shock transcription factors in plant heat stress tolerance: a brief appraisal’, Molecular Biology Reports, 49(6), pp. 5771– 5785. Available at: 10.1007/s11033-022-07190-x.

Hasan, Md.M., et al. (2021) ‘GABA: A Key Player in Drought Stress Resistance in Plants’, International Journal of Molecular Sciences, 22(18), p. 10136. Available at: 10.3390/ijms221810136.

Hilker, M., et al. (2016) ‘Priming and memory of stress responses in organisms lacking a nervous system’, Biological Reviews, 91(4), pp. 1118–1133. Available at: 10.1111/brv.12215.

Ho, C.-H., et al. (2009) ‘CHL1 Functions as a Nitrate Sensor in Plants’, Cell, 138(6), pp. 1184–1194. Available at: 10.1016/j.cell.2009.07.004.

Hu, Y., Dong, Q. and Yu, D. (2012) ‘*Arabidopsis* WRKY46 coordinates with WRKY70 and WRKY53 in basal resistance against pathogen *Pseudomonas syringae*’, Plant Science, 185–186, pp. 288–297. Available at: 10.1016/j.plantsci.2011.12.003.

Islam, M.J., et al. (2022) ‘Exogenous putrescine attenuates the negative impact of drought stress by modulating physio-biochemical traits and gene expression in sugar beet (Beta vulgaris L.)’, PLoS ONE, 17(1), p. e0262099. Available at: 10.1371/journal.pone.0262099.

du Jardin, P. (2015) ‘Plant biostimulants: Definition, concept, main categories and regulation’, Scientia Horticulturae, 196, pp. 3–14. Available at: 10.1016/j.scienta.2015.09.021.

Kanehisa, M. and Goto, S. (2000) ‘KEGG: Kyoto Encyclopedia of Genes and Genomes’, Nucleic Acids Research, 28(1), pp. 27–30. Available at: 10.1093/nar/28.1.27.

Kawasaki, S., et al. (2000) ‘Responses of Wild Watermelon to Drought Stress: Accumulation of an ArgE Homologue and Citrulline in Leaves during Water Deficits’, Plant and Cell Physiology, 41(7), pp. 864–873. Available at: 10.1093/pcp/pcd005.

Khan, A., et al. (2019) ‘Silicon and salicylic acid confer high-pH stress tolerance in tomato seedlings’, Scientific Reports, 9(1), p. 19788. Available at: 10.1038/s41598-019-55651-4.

Kotak, S., et al. (2007) ‘Complexity of the heat stress response in plants’, Current Opinion in Plant Biology, 10(3), pp. 310–316. Available at: 10.1016/j.pbi.2007.04.011.

Le S., Josse, J. and Husson, F. (2008) ‘FactoMineR: An R Package for Multivariate Analysis’, Journal of Statistical Software, 25, pp. 1–18. Available at: 10.18637/jss.v025.i01.

Lei, S. and Huang, B. (2022) ‘Metabolic regulation of α-Ketoglutarate associated with heat tolerance in perennial ryegrass’, Plant Physiology and Biochemistry, 190, pp. 164–173. Available at: 10.1016/j.plaphy.2022.09.005.

Li, A., Sun, X. and Liu, L. (2022) ‘Action of Salicylic Acid on Plant Growth’, Frontiers in Plant Science, 13. Available at: 10.3389/fpls.2022.878076.

Li, J., et al. (2015) ‘SHOEBOX Modulates Root Meristem Size in Rice through Dose-Dependent Effects of Gibberellins on Cell Elongation and Proliferation’, PLOS Genetics, 11(8), p. e1005464. Available at: 10.1371/journal.pgen.1005464.

Li, J.-Y., et al. (2010a) ‘The Arabidopsis Nitrate Transporter NRT1.8 Functions in Nitrate Removal from the Xylem Sap and Mediates Cadmium Tolerance’, The Plant Cell, 22(5), pp. 1633–1646. Available at: 10.1105/tpc.110.075242.

Li, J.-Y., et al. (2010b) ‘The Arabidopsis Nitrate Transporter NRT1.8 Functions in Nitrate Removal from the Xylem Sap and Mediates Cadmium Tolerance’, The Plant Cell, 22(5), pp. 1633–1646. Available at: 10.1105/tpc.110.075242.

Li, Y., Jones, L. and McQueen-Mason, S. (2003) ‘Expansins and cell growth’, Current Opinion in Plant Biology, 6(6), pp. 603–610. Available at: 10.1016/j.pbi.2003.09.003.

Livanos, P., Apostolakos, P. and Galatis, B. (2012) ‘Plant cell division: ROS homeostasis is required’, Plant Signaling & Behavior, 7(7), pp. 771–778. Available at: 10.4161/psb.20530.

Lu, Y., et al. (2016) ‘Genome-wide identification and expression analysis of the expansin gene family in tomato’, Molecular Genetics and Genomics, 291(2), pp. 597–608. Available at: 10.1007/s00438-015-1133-4.

Malécange, M., et al. (2022) ‘Leafamine®, a Free Amino Acid-Rich Biostimulant, Promotes Growth Performance of Deficit-Irrigated Lettuce’, International Journal of Molecular Sciences, 23(13), p. 7338. Available at: 10.3390/ijms23137338.

Malécange, M., et al. (2023) ‘Biostimulant Properties of Protein Hydrolysates: Recent Advances and Future Challenges’, International Journal of Molecular Sciences, 24(11), p. 9714. Available at: 10.3390/ijms24119714.

Mauch-Mani, B., et al. (2017) ‘Defense Priming: An Adaptive Part of Induced Resistance’, Annual Review of Plant Biology, 68(1), pp. 485–512. Available at: 10.1146/annurev-arplant-042916-041132.

Mesihovic, A., et al. (2022) ‘HsfA7 coordinates the transition from mild to strong heat stress response by controlling the activity of the master regulator HsfA1a in tomato’, Cell Reports, 38(2). Available at: 10.1016/j.celrep.2021.110224.

Morales de los Ríos, L., et al. (2021) ‘The Arabidopsis protein NPF6.2/NRT1.4 is a plasma membrane nitrate transporter and a target of protein kinase CIPK23’, Plant Physiology and Biochemistry, 168, pp. 239–251. Available at: 10.1016/j.plaphy.2021.10.016.

Moussu, S. and Ingram, G. (2023) ‘The EXTENSIN enigma’, The Cell Surface, 9, p. 100094. Available at: 10.1016/j.tcsw.2023.100094.

Naulin, P.A., et al. (2020) ‘Nitrate Induction of Primary Root Growth Requires Cytokinin Signaling in Arabidopsis thaliana’, Plant and Cell Physiology, 61(2), pp. 342–352. Available at: 10.1093/pcp/pcz199.

Ninkuu, V., et al. (2025) ‘Phenylpropanoids metabolism: recent insight into stress tolerance and plant development cues’, Frontiers in Plant Science, 16. Available at: 10.3389/fpls.2025.1571825.

Niu, Y. and Xiang, Y. (2018) ‘An Overview of Biomembrane Functions in Plant Responses to High-Temperature Stress’, Frontiers in Plant Science, 9. Available at: 10.3389/fpls.2018.00915.

Nogueira, M., et al. (2024) ‘Ketocarotenoid production in tomato triggers metabolic reprogramming and cellular adaptation: The quest for homeostasis’, Plant Biotechnology Journal, 22(2), pp. 427–444. Available at: 10.1111/pbi.14196.

Pasternak, T., et al. (2019) ‘Salicylic Acid Affects Root Meristem Patterning via Auxin Distribution in a Concentration-Dependent Manner’, Plant Physiology, 180(3), pp. 1725– 1739. Available at: 10.1104/pp.19.00130.

Peng, Y., et al. (2021) ‘Salicylic Acid: Biosynthesis and Signaling’, Annual Review of Plant Biology, 72(Volume 72, 2021), pp. 761–791. Available at: 10.1146/annurev-arplant-081320-092855.

Potuschak, T. and Doerner, P. (2001) ‘Cell cycle controls: genome-wide analysis in Arabidopsis’, Current Opinion in Plant Biology, 4(6), pp. 501–506. Available at: 10.1016/S1369-5266(00)00207-7.

Pratt, W.B., et al. (2010) ‘Role of the Hsp90/Hsp70-Based Chaperone Machinery in Making Triage Decisions When Proteins Undergo Oxidative and Toxic Damage’, Experimental biology and medicine (Maywood, N.J.), 235(3), pp. 278–289. Available at: 10.1258/ebm.2009.009250.

Riou-Khamlichi, C., et al. (1999) ‘Cytokinin Activation of Arabidopsis Cell Division Through a D-Type Cyclin’, Science, 283(5407), pp. 1541–1544. Available at: 10.1126/science.283.5407.1541.

Samalova, M., et al. (2020) ‘Expansin-controlled cell wall stiffness regulates root growth in Arabidopsis’. bioRxiv, p. 2020.06.25.170969. Available at: 10.1101/2020.06.25.170969.

Sampedro, J. and Cosgrove, D.J. (2005) ‘The expansin superfamily’, Genome Biology, 6(12), p. 242. Available at: 10.1186/gb-2005-6-12-242.

Sato, S., Peet, M.M. and Thomas, J.F. (2000) ‘Physiological factors limit fruit set of tomato (Lycopersicon esculentum Mill.) under chronic, mild heat stress’, Plant, Cell & Environment, 23(7), pp. 719–726. Available at: 10.1046/j.1365-3040.2000.00589.x.

Sestili, F. et al. (2018) ‘Protein Hydrolysate Stimulates Growth in Tomato Coupled With N-Dependent Gene Expression Involved in N Assimilation’, Frontiers in Plant Science, 9. Available at: https://www.frontiersin.org/articles/10.3389/fpls.2018.01233 (Accessed: 15 August 2022).

Shah Jahan, M., et al. (2019) ‘Exogenous salicylic acid increases the heat tolerance in Tomato (*Solanum lycopersicum* L) by enhancing photosynthesis efficiency and improving antioxidant defense system through scavenging of reactive oxygen species’, Scientia Horticulturae, 247, pp. 421–429. Available at: 10.1016/j.scienta.2018.12.047.

Song, Q., et al. (2020) ‘Functional Relevance of Citrulline in the Vegetative Tissues of Watermelon During Abiotic Stresses’, Frontiers in Plant Science, 11. Available at: 10.3389/fpls.2020.00512.

Suliman, A.A., et al. (2024) ‘Boosting Resilience and Efficiency of Tomato Fields to Heat Stress Tolerance Using Cytokinin (6-Benzylaminopurine)’, Horticulturae, 10(2), p. 170. Available at: 10.3390/horticulturae10020170.

Sun, L., et al. (2015) ‘Candidate gene selection and detailed morphological evaluations of fs8.1, a quantitative trait locus controlling tomato fruit shape’, Journal of Experimental Botany, 66(20), pp. 6471–6482. Available at: 10.1093/jxb/erv361.

Tegeder, M. and Ward, J.M. (2012) ‘Molecular Evolution of Plant AAP and LHT Amino Acid Transporters’, Frontiers in plant science, 3, p. 21. Available at: 10.3389/fpls.2012.00021.

Teixeira, W.F., et al. (2017) ‘Foliar and Seed Application of Amino Acids Affects the Antioxidant Metabolism of the Soybean Crop’, Frontiers in Plant Science, 8. Available at: 10.3389/fpls.2017.00327.

Tiwari, M., et al. (2022) ‘Genetic and molecular mechanisms underlying root architecture and function under heat stress—A hidden story’, *Plant*, Cell & Environment, 45(3), pp. 771–788. Available at: 10.1111/pce.14266.

Vaseva, I.I., et al. (2022) ‘Heat-Stress-Mitigating Effects of a Protein-Hydrolysate-Based Biostimulant Are Linked to Changes in Protease, DHN, and HSP Gene Expression in Maize’, Agronomy, 12(5), p. 1127. Available at: 10.3390/agronomy12051127.

Verma, V., Ravindran, P. and Kumar, P.P. (2016) ‘Plant hormone-mediated regulation of stress responses’, BMC Plant Biology, 16(1), p. 86. Available at: 10.1186/s12870-016-0771-y.

Wang, W., et al. (2004) ‘Role of plant heat-shock proteins and molecular chaperones in the abiotic stress response’, Trends in Plant Science, 9(5), pp. 244–252. Available at: 10.1016/j.tplants.2004.03.006.

Wang, W., et al. (2022) ‘Animal-derived plant biostimulant alleviates drought stress by regulating photosynthesis, osmotic adjustment, and antioxidant systems in tomato plants’, Scientia Horticulturae, 305, p. 111365. Available at: 10.1016/j.scienta.2022.111365.

Wu, H.-C., Bulgakov, V.P. and Jinn, T.-L. (2018) ‘Pectin Methylesterases: Cell Wall Remodeling Proteins Are Required for Plant Response to Heat Stress’, Frontiers in Plant Science, 9. Available at: 10.3389/fpls.2018.01612.

Wu, T., et al. (2021) ‘clusterProfiler 4.0: A universal enrichment tool for interpreting omics data’, The Innovation, 2(3), p. 100141. Available at: 10.1016/j.xinn.2021.100141.

Xu, P., et al. (2014) ‘HDG11 upregulates cell-wall-loosening protein genes to promote root elongation in Arabidopsis’, Journal of Experimental Botany, 65(15), pp. 4285–4295. Available at: 10.1093/jxb/eru202.

Yakhin, O.I. et al. (2017) ‘Biostimulants in Plant Science: A Global Perspective’, Frontiers in Plant Science, 7. Available at: https://www.frontiersin.org/articles/10.3389/fpls.2016.02049 (Accessed: 15 August 2022).

Yang, X., et al. (2016) ‘Heat shock factors in tomatoes: genome-wide identification, phylogenetic analysis and expression profiling under development and heat stress’, PeerJ, 4, p. e1961. Available at: 10.7717/peerj.1961.

Zhao, J., et al. (2021) ‘Exogenous Putrescine Alleviates Drought Stress by Altering Reactive Oxygen Species Scavenging and Biosynthesis of Polyamines in the Seedlings of Cabernet Sauvignon’, Frontiers in Plant Science, 12. Available at: 10.3389/fpls.2021.767992.

