## Supplemental Figure 1 for "The free amino acid-rich biostimulant, Leafamine®, promotes cell division in tomato roots and alleviates heat stress effects"

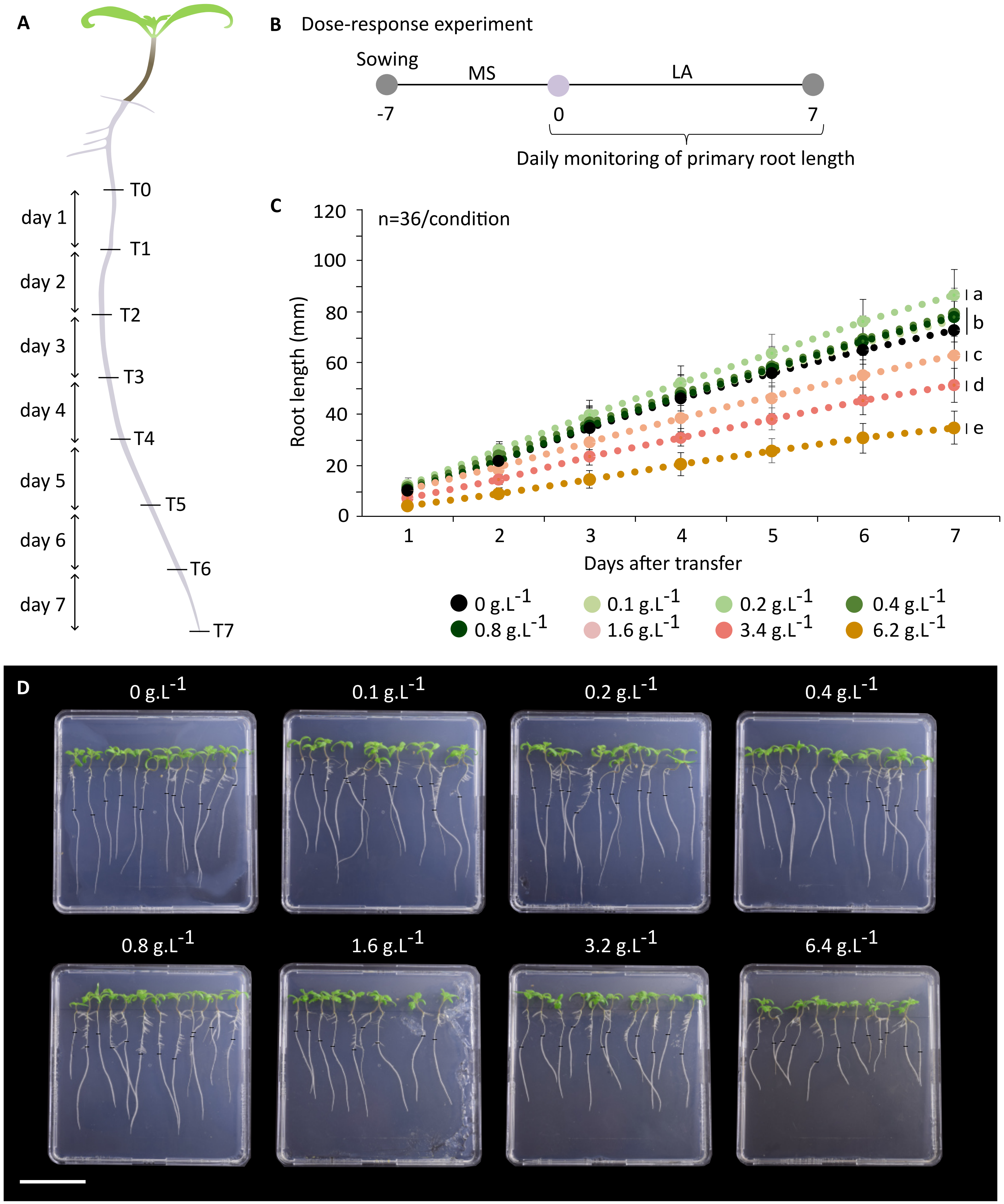


**Figure S1.** Dose response of LA treatment on primary root development of tomato seedlings. (A) Schematic representation of a tomato plantlet and the time points for root length measurement. (B) Experimental design of the dose-response assay for LA treatment. (C) Primary root growth over time in seedlings treated with different LA concentrations. (D) Representative images of plates with seedlings grown on media supplemented with increasing concentrations of LA. Scale bar = 8 cm. Statistical comparisons (p-value) are shown in Table S2. Letters indicate significant differences between treatments. ANOVA test was used followed by Tukey's multiple comparisons test.
