## Supplemental Figure 2 for "The free amino acid-rich biostimulant, Leafamine®, promotes cell division in tomato roots and alleviates heat stress effects"

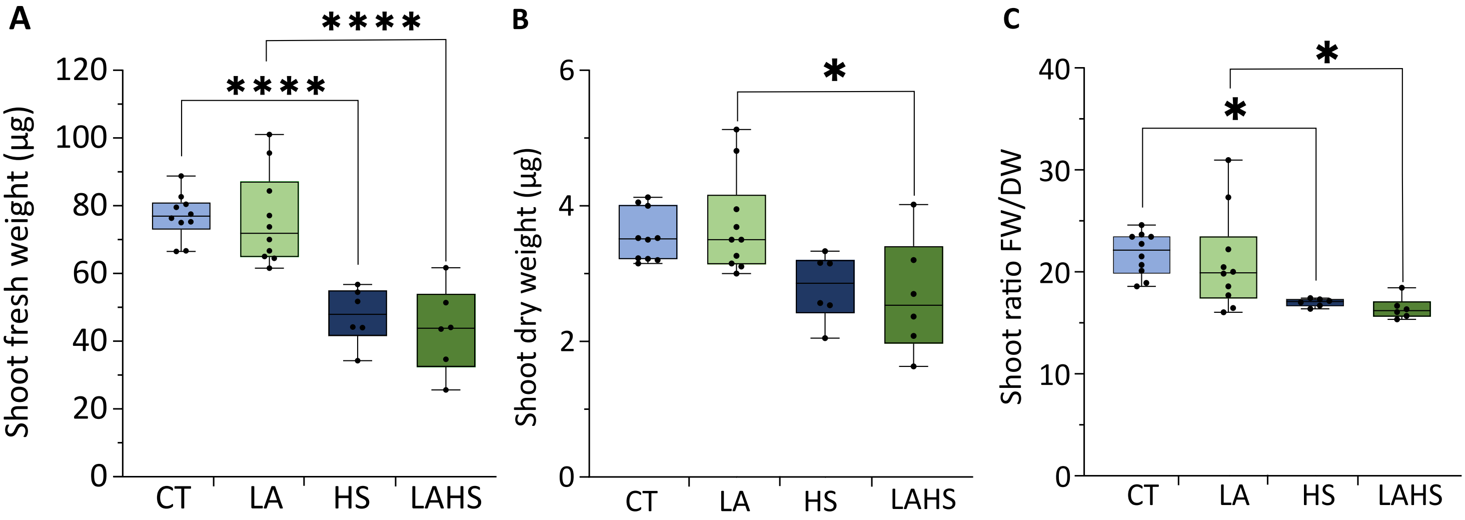


**Figure S2.** Effect of the LA treatment on shoot growth of tomato seedlings grown under NS and heat stress HS conditions for 7 days. (A) Shoot fresh weight. (B) Shoot dry weight. (C) Shoot fresh-to-dry weight ratio (FW/DW). Boxplot: whiskers extend from minimum to maximum, box extends 25th to 75th percentiles, the line in the middle is the median. Letters indicate significant differences among treatments. Asterisks indicate significant differences. ANOVA test was used followed by Tukey's multiple comparisons test; *$p < 0.05$, $p < 0.005$, *$p < 0.0005$, ****$p < 0.0001$.
