## Supplemental Figure 3 for "The free amino acid-rich biostimulant, Leafamine®, promotes cell division in tomato roots and alleviates heat stress effects"

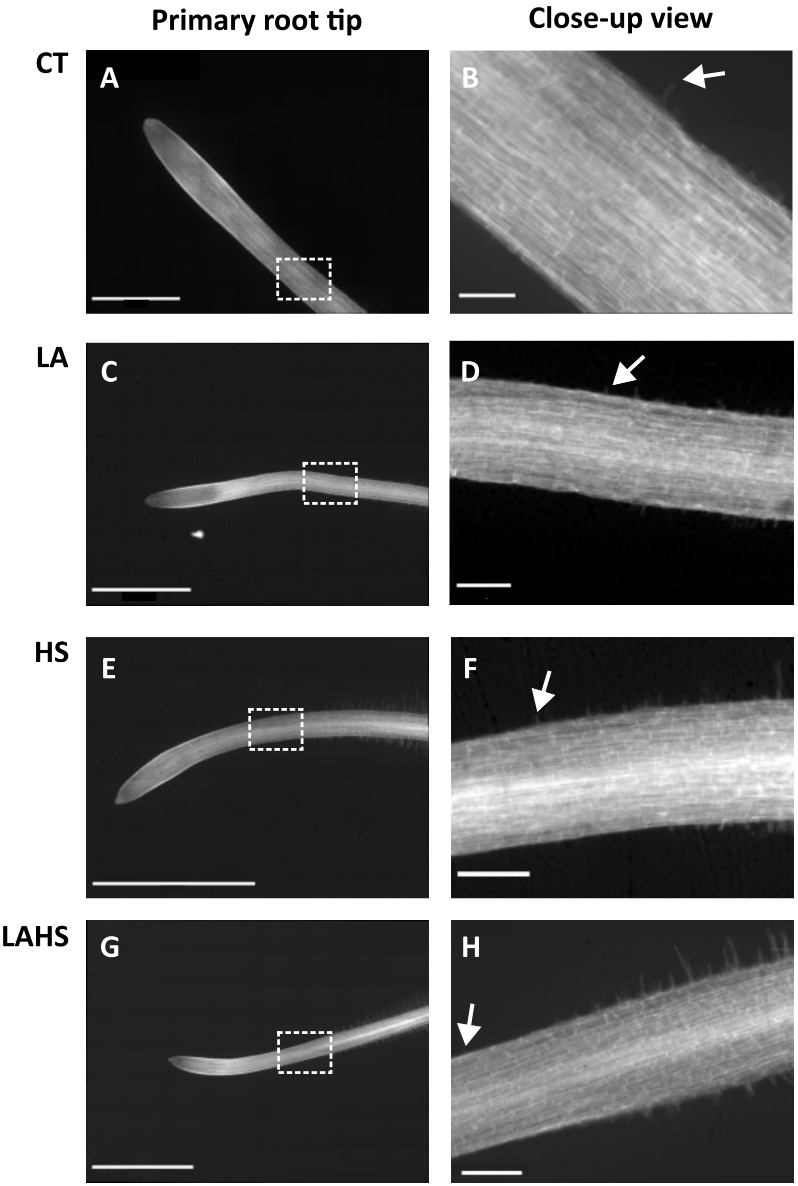


**Figure S3.** Effect of LA on the length of the elongation zone of the primary root of seedlings grown under NS and HS conditions. (A, C, E, G) Primary root tip of untreated, unstressed plants (CT) (A); LA-treated, unstressed plants (LA) (C); untreated, heat-stressed plants (HS) (E); LA-treated, heat stressed plant (LAHS) (G). Scale bar = 2mm. (B, D, F, H) Close-up view (white rectangle in A, C, E, G) of the region presenting the upper boundary of the elongation zone for each condition, defined as the position of the first root hair (white Arrows). Scale bar = 200µm
