## Supplemental Figure 4 for "The free amino acid-rich biostimulant, Leafamine®, promotes cell division in tomato roots and alleviates heat stress effects"

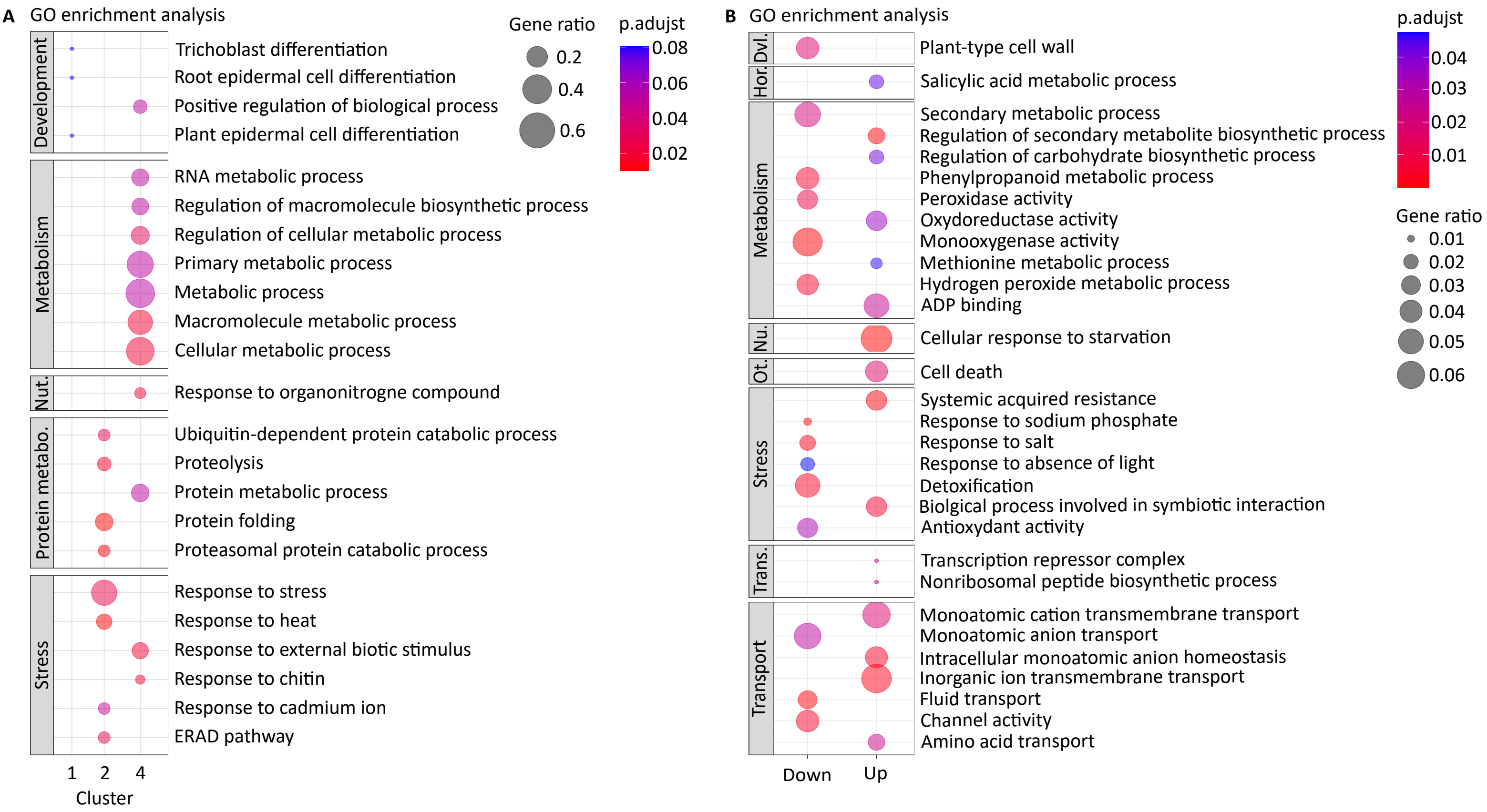


**Figure S4.** LA treatment exerts condition-dependent effects under HS, specifically impacting molecular processes related to metabolism and stress responses. (A) GO enrichment analysis of each cluster resulting from the LA x HS interaction. (B) GO enrichment analysis of DEGs (Log2FC > 0.5) consistently up- or downregulated by LA treatment, under both NS and HS conditions. Dvl = Development. Hor = Hormones. Metabo = Metabolism. Nu = Nutrition. Ot = Others. Trans = Transcription. GO terms are grouped by category; dot color represents adjusted p-values, and dot size indicates gene ratio. GO terms from table S4, table S5, and table S6 were grouped to remove redundancy.
