## Supplemental Figure 5 for "The free amino acid-rich biostimulant, Leafamine®, promotes cell division in tomato roots and alleviates heat stress effects"

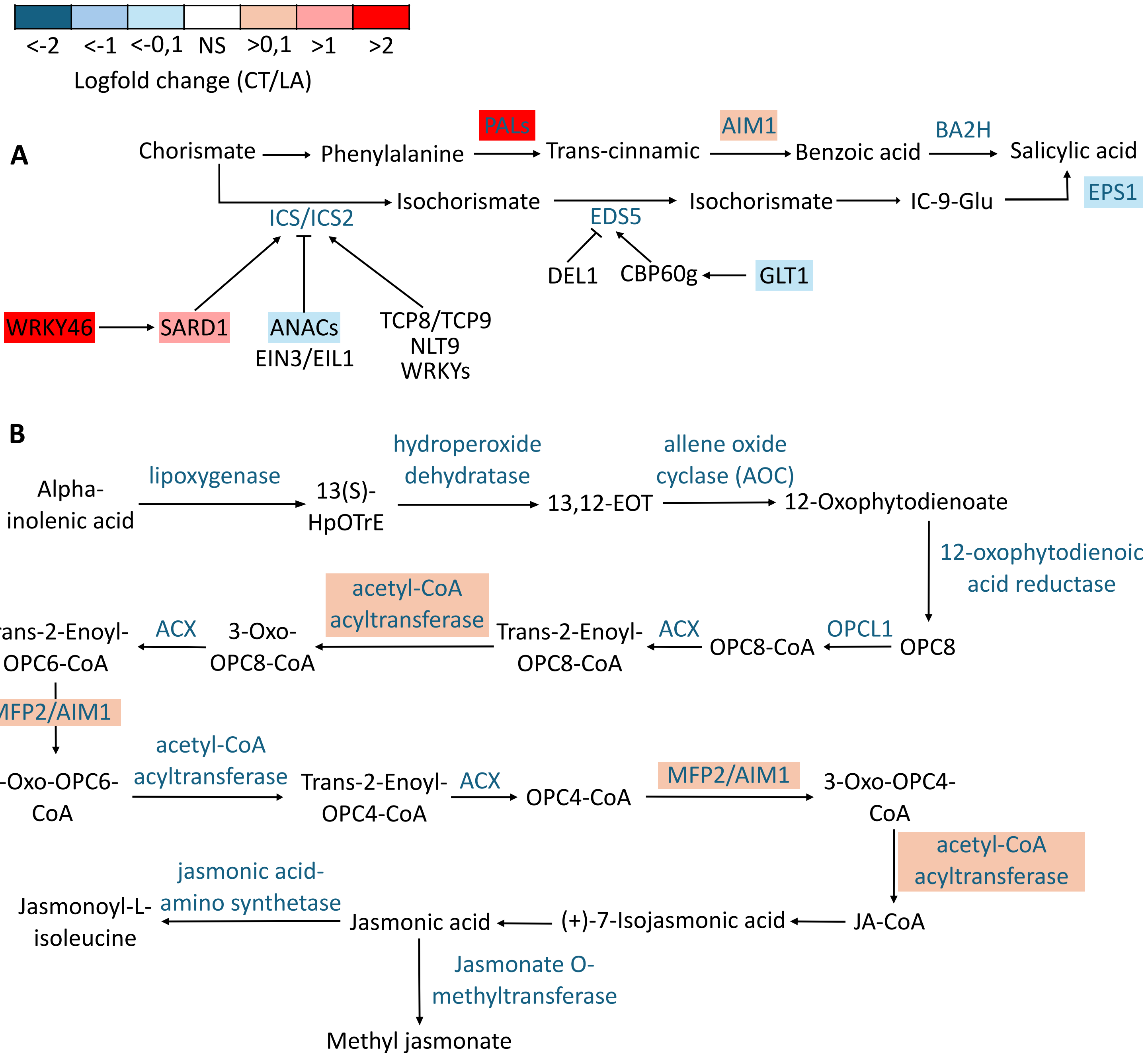


**Figure S5.** Schemes representing the transcriptional changes of phytohormone biosynthesis pathways in roots five days after LA treatment under NS conditions based on the transcriptomic data. (A) Expression changes in genes involved in the SA biosynthetic pathway. (B) Expression changes in genes involved in the JA biosynthetic pathway. The schematic shows key enzymes and intermediates of each pathway. Colour shading represents the $log_2$ fold-change ratio (CT/LA) in gene expression, with red indicating upregulation and blue indicating downregulation. Genes with statistically significant expression changes ($p < 0.05$) are marked by coloured boxes. The colour scale bar indicates the magnitude and direction of the change.
